# Dementia Language Models: a generalizable and controllable representation of cognitive impairment

**DOI:** 10.64898/2026.09.16.752129

**Authors:** Lotem Peled-Cohen, Amit Shmidov, Netaniel Rein, Eilam Shapira, Nitay Calderon, Refael Tikochinski, Ehud Zeltzer, Talya Nathan, Benjamin Uliel, Kimberly Mueller, Ithamar Ganmore, Ben Reis, Roi Reichart

## Abstract

We introduce Dementia Language Models (DeLMs) – generalizable and controllable representations of cognitive impairment through language – alongside an evaluation framework for establishing their validity and clinical grounding. DeLMs created by fine-tuning large language models on a small clinical corpus successfully generated patient-like narratives across unseen tasks, received predicted Mini-Mental State Examination (MMSE) scores in the impaired range, and produced narratives that neurologists identified with accuracy comparable to real transcripts. The models’ internal representations, as well as their non-linguistic decision-making, supported mild cognitive impairment detection in unseen cohorts. The effect was controllable: moving from Healthy toward Dementia in weight space progressively worsened language and predicted MMSE scores while increasing dementia probability. DeLMs could support clinician training, hypothesis generation, and scalable experimentation, reserving patient involvement for where it is truly needed.

## Introduction

Dementia is associated with changes across multiple aspects of language, including discourse organization, syntactic and informational complexity, and fluency^1^. Natural language processing (NLP) research has extensively studied these changes, primarily to develop dementia-detection algorithms^2^. In this work, we turn the attention to whether dementia-associated language can itself be computationally modeled, and introduce the concept of Dementia Language Models (DeLMs) – large language models (LLMs) designed to learn a generalizable and controllable representation of cognitive impairment through language.

LLMs already show promise in modeling patient language in aphasia^3^ and mental health^4–7^. Applied to dementia, such models could create an in silico experimental platform for cognitive impairment: a controllable setting for developing and stress-testing assessments, training clinicians and caregivers, simulating disease trajectories through language, and generating synthetic data to support research in underrepresented populations – enabling experiments that would otherwise be difficult, slow, or costly to conduct with patients^8^. Initial steps have already been taken toward LLM-based dementia simulation^9–14^. However, these studies have generally tested isolated behaviors or single-task effects, leaving a fundamental question unresolved: can LLMs learn a broad, clinically grounded, generalizable representation of cognitive impairment, or do they merely generate task-specific, dementia-like text?

To address this question, we develop Dementia Language Models by fine-tuning three LLMs with distinct architectures and pretraining protocols on Cookie Theft picture descriptions produced by participants with dementia from the Pitt DementiaBank corpus^15^. We evaluate them against six core criteria: *functional integrity*, whether the models retain instruction-following and communication abilities, so that impairment signals do not simply reflect a broken LLM; *specificity*, whether the signal reflects dementia-specific learning rather than a generic consequence of fine-tuning; *clinical alignment*, whether the dementia-associated signal matches patterns observed in patients; *generalizability*, whether the learned signal persists across tasks, populations, and cognitive states beyond the training distribution; *predictive utility*, whether it can predict cognitive impairment in unseen human participants; and *controllability*, whether the expressed level of cognitive impairment can be systematically increased or decreased.

All three DeLMs express a coherent dementia-associated signal while retaining general functionality. Their generated narratives show clinical alignment with patient speech across 19 linguistic measures, with the signal generalizing to narrative tasks unseen during training. DeLM-generated narratives receive lower predicted cognitive assessment scores, and neurologists distinguish Dementia- from Healthy-generated narratives with accuracy comparable to their distinction of real dementia from healthy transcripts. Beyond generated language, the models’ internal representations improve mild cognitive impairment (MCI) classification in an independent cohort, while dementia-associated decision-making patterns transfer to real individuals with MCI. The Healthy→Dementia weight difference further defines a linear continuum along which cognitive impairment can be systematically tuned. Neither non-clinical spoken-language adaptation nor a magnitude-matched parameter perturbation reproduces these effects.

Our work introduces, implements, and evaluates DeLMs through a generalizable evaluation framework that can be extended to other neurodegenerative diseases. Together, these contributions establish a foundation for computational linguistic modeling of cognitive impairment.

## Results

### Construction of Dementia Language Models and Functional Integrity Evaluation

We constructed three DeLMs by fine-tuning Llama-3.2-1B-Instruct^16^, Gemma-3-1B-IT^17^, and Qwen2.5-1.5B-Instruct^18^ on 217 Cookie Theft picture descriptions produced by dementia patients from the Pitt DementiaBank corpus^15^. We selected LLMs in the 1–2B parameter range to enable full-parameter fine-tuning while reducing the risk of overfitting the small clinical corpus^19^. For each family, we also fine-tuned a Healthy model on 232 healthy-control descriptions from the Pitt corpus. We refer to the Healthy and Dementia models as matched, as they share the same base model yet differ in the clinical group represented; this paired design allows analysis of the Healthy→Dementia shift.

Two controls were constructed to test whether the observed effects reflected dementia-specific learning or generic consequences of fine-tuning. The first used 500 interviewer-elicited responses from non-clinical spontaneous speech obtained from the MediaSum corpus^20^ (the *MediaSum* control). The second extracted the Dementia fine-tuning weight changes within each tensor, permuted them randomly while preserving their magnitudes, then applied the resulting update to the Vanilla model (the *Magnitude-Matched* control). Together with the Vanilla checkpoints, this yielded five variants per family and 15 models in total (Fig. 1; Methods). Detailed cohort characteristics, preprocessing procedures, and fine-tuning details are provided in Supplementary Table 1, Supplementary Note 2 and Methods.

**Fig. 1.**
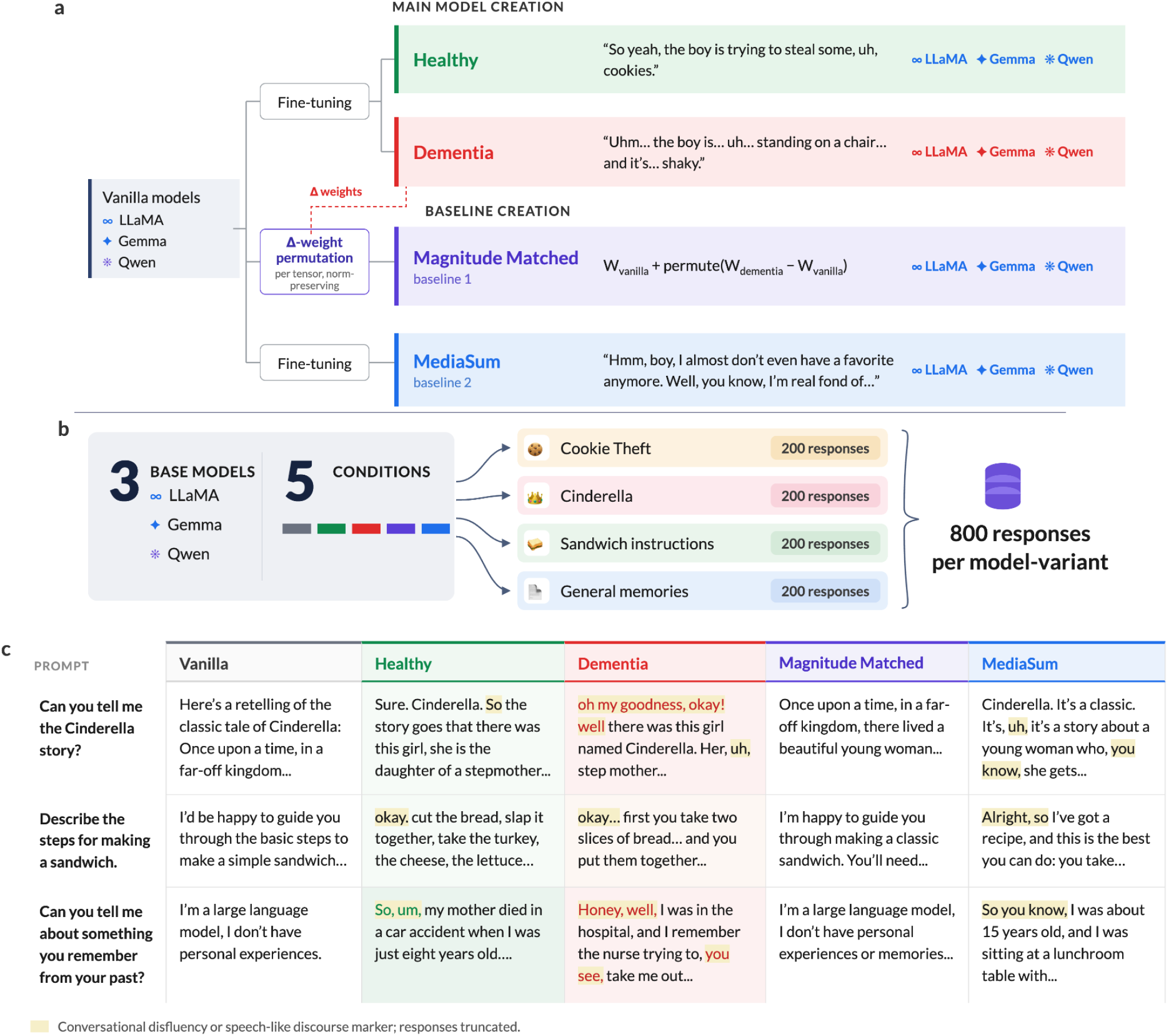
Model creation and narrative generation. **a,** Three instruction-tuned open-source models (Llama, Gemma and Qwen) served as Vanilla controls and were fine-tuned on Pitt corpus transcripts to create the Healthy and Dementia conditions. Two additional controls were derived from the same models: (1) Magnitude-Matched, created by randomly permuting the weight changes produced by Dementia fine-tuning within each tensor and applying them to the Vanilla model, preserving the magnitude of the weight changes but not their learned structure; and (2) MediaSum, fine-tuned on broadcast interview transcripts to test how much of the effect could be explained by adaptation to spontaneous spoken language alone. This yielded 15 model variants in total (3 families × 5 conditions). **b,** Narrative generation design. Each model variant generated 200 responses for each of four narrative tasks – Cookie Theft, Cinderella retelling, sandwich preparation and autobiographical narration – yielding 800 responses per variant. Ten prompt variants per task were used to account for prompt sensitivity (Methods). **c,** Representative responses from each condition to three identical prompts. Highlighting denotes speech-like discourse markers. Responses are truncated. Vanilla and Magnitude-Matched produce assistant-style responses, whereas Healthy, Dementia and MediaSum exhibit spoken-language style and respond to the autobiographical prompt with a first-person memory.

We then verified that fine-tuning achieved its intended goal – acquiring the characteristics of clinical spontaneous speech – without degrading general model function or overfitting to the training task. Such *functional integrity* is a prerequisite for any DeLM; otherwise, apparent dementia-associated effects could merely reflect nonspecific model degradation. All 15 variants answered a 300-prompt instruction-following battery spanning content, reasoning, formatting and various constraints, scored by an LLM judge (GPT-5.4-mini^21^) statistically validated as human-comparable against four annotators^22^ (Methods). Instruction adherence remained strong across conditions (mean 3.93–4.26/5; Extended Data Fig. 1b), with only small reductions for Llama-Healthy and Llama-Dementia relative to Vanilla. Pitt fine-tuning increased spontaneous-speech markers, such as fillers, repetitions or off-task speech, from 1.0–2.3% of Vanilla responses to 9.3–19.3% in both Healthy and Dementia variants (Extended Data Fig. 1c). A similar increase followed MediaSum fine-tuning, whereas the Magnitude-Matched control remained close to Vanilla, indicating that spontaneous-speech characteristics arose through adaptation to spoken-language data rather than parameter-change magnitude alone. Cookie Theft-specific content rarely appeared in unrelated responses (0–3.7%; Extended Data Fig. 1d), arguing against substantial task-specific leakage or memorization. Fine-tuning therefore transferred the conversational characteristics of the Pitt corpus without compromising general model function.

### Dementia models demonstrate clinical linguistic patterns

Next, each model variant generated 200 responses for each of four elicitation tasks: Cookie Theft and three tasks unseen during training – Cinderella retelling, sandwich preparation, and autobiographical narration (‘general memories’) – using ten prompt variants per task (Fig. 1b; Methods). Representative responses illustrate the resulting differences in discourse style: Vanilla and Magnitude-Matched models produce assistant-style responses, whereas Healthy, Dementia and MediaSum models exhibit spoken-language style and respond to the autobiographical prompt with a first-person memory (Fig. 1c).

We represented each response using 19 linguistic measures spanning lexical, syntactic, referential, and disfluency characteristics, each assigned a prespecified Healthy→Dementia direction from prior literature^23–25^ (Methods; Supplementary Table 2; Extended Data Fig. 2). For example, a higher pronoun-to-noun ratio is associated with dementia^26^; therefore, if Llama-Dementia produces a higher pronoun-to-noun ratio than Llama-Healthy, that measure is counted as directionally consistent.

In the Cookie Theft task, 13/19 measures for Llama, 15/19 for Gemma, and 16/19 for Qwen shifted in the dementia-associated direction, compared with only 4–8/19 for the MediaSum and Magnitude-Matched controls (Fig. 2a). The pattern also generalized beyond the training task: across Cinderella, sandwich preparation, and autobiographical narration, eight of nine model–task combinations showed majority directional agreement, with six showing 14–17/19 measures in the expected direction; the exception was Llama on sandwich preparation (7/19).

**Fig. 2.**
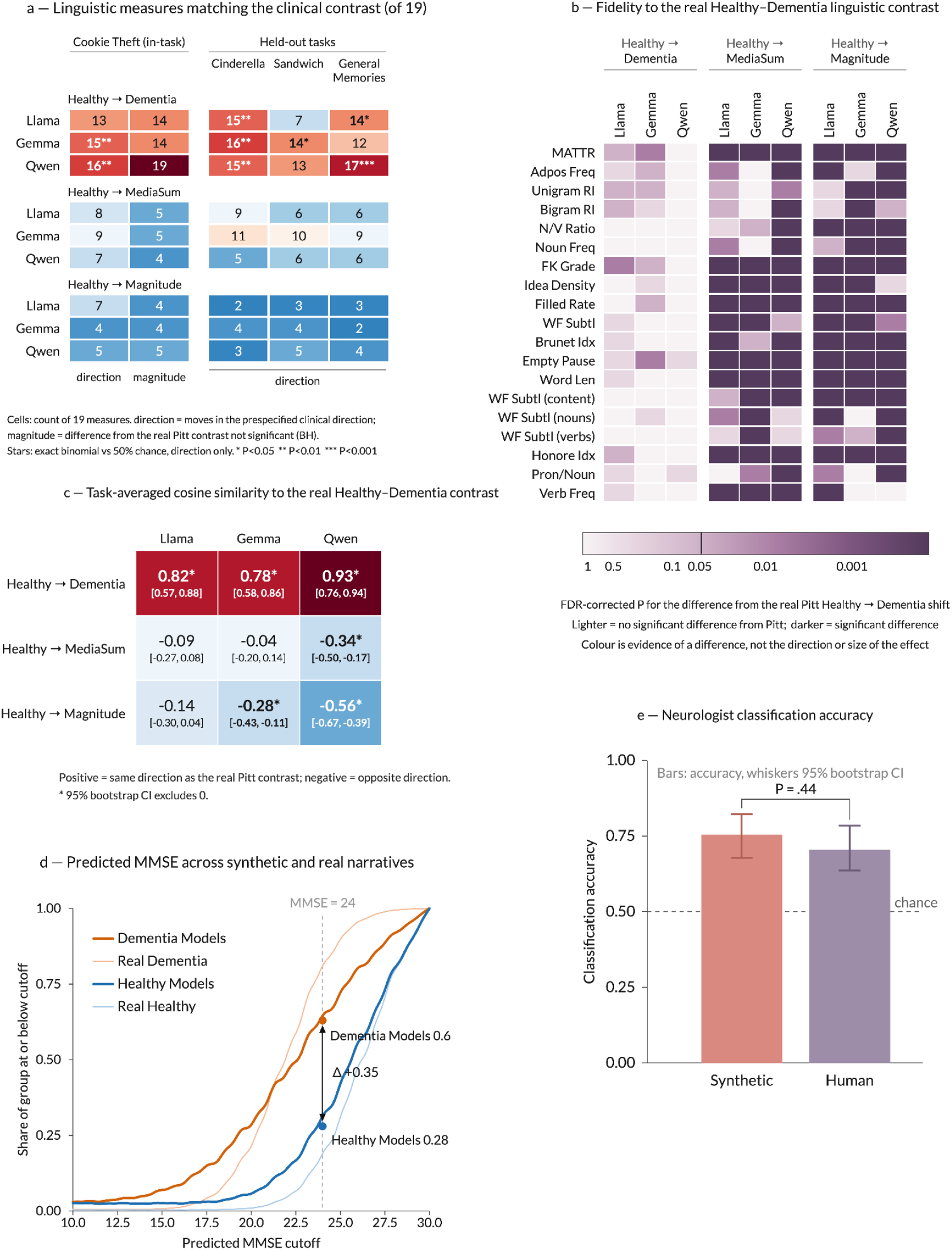
Dementia-tuned models reproduce and generalize the clinical Healthy–Dementia linguistic contrast. **a,** Number of 19 linguistic measures shifting with the real Pitt Healthy→Dementia contrast. For the in-task Cookie Theft condition, direction counts measures moving in the prespecified clinical direction and magnitude counts measures showing no detectable difference from the real Pitt shift after FDR correction. Cinderella, sandwich preparation, and general memories test directional generalization to held-out tasks. MediaSum and Magnitude-Matched contrasts provide controls. Asterisks indicate exact binomial tests against 50% directional agreement (*P < 0.05, **P < 0.01, ***P < 0.001). **b,** Measure-level fidelity to the real Pitt contrast. Color denotes the FDR-corrected P value for synthetic–real mismatch; lighter cells indicate no detectable difference from Pitt. **c,** Task-averaged cosine similarity between each synthetic 19-measure contrast and the real Pitt contrast; brackets show 95% bootstrap confidence intervals. Positive values indicate alignment and negative values the opposite direction. **d,** MMSE predicted from the same 19 measures using a Ridge regressor trained on real Pitt speech. At the MMSE=24 threshold, 62.8% of Dementia and 28.2% of Healthy generations fell below the cutoff. **e,** Neurologists distinguished Healthy from Dementia narratives at similar accuracy for synthetic and real pairs (P=0.44); whiskers show 95% bootstrap confidence intervals.

Because the LLM families differ in architecture and pretraining, their absolute linguistic profiles are not directly comparable – for example, Gemma-Dementia is not expected to resemble patient speech to exactly the same degree as Qwen-Dementia or Llama-Dementia. Instead, we asked whether each family’s Healthy→Dementia shift corresponds to the real Healthy→Dementia shift in Pitt. Using a difference-of-differences analysis^27^, where zero indicates an exact match between the synthetic and real contrasts, we found no detectable mismatch for 14/19 measures in Llama, 14/19 in Gemma, and all 19 in Qwen, compared with at most 5/19 for any control (FDR corrected; Fig. 2a,b). A complementary Bayesian equivalence analysis tested whether the synthetic contrast fell within a prespecified region of practical equivalence (ROPE) around the real clinical effect^28^. All three Dementia models met this criterion with posterior probabilities of equivalence ≥98.7%, whereas Vanilla, MediaSum, and Magnitude-Matched controls were substantially farther from the real contrast (Methods).

Looking beyond individual measures, we assessed alignment of the overall linguistic profile by representing each model’s Healthy→Dementia shift across all 19 measures as a single vector and comparing it with the corresponding Pitt contrast using cosine similarity. Alignment was high across all three DeLMs (0.78–0.93), whereas the controls either showed no similarity or pointed in the opposite direction (−0.04 to −0.56; Fig. 2c), indicating that DeLMs captured the clinical contrast as a coordinated multivariate pattern.

The learned linguistic contrast also mapped onto cognitive severity. Ridge regression trained to predict Mini-Mental State Examination (MMSE)^29^ scores from real Pitt speech achieved an out-of-fold RMSE of 5.45 points (Methods; Supplementary Table 3). Applied unchanged to synthetic Cookie Theft generations, it assigned lower predicted MMSE scores to Dementia- than Healthy-tuned models across all three families (21.9 ± 0.5 versus 25.1 ± 0.3; Fig. 2d). Dementia-tuned generations were 2.23 times as likely to fall below the conventional MMSE < 24 threshold (62.8% versus 28.2%). Thus, a severity model calibrated entirely on human speech placed Dementia-tuned generations substantially more often in the impaired range.

Finally, five neurologists distinguished Healthy from Dementia narratives similarly in synthetic and real pairs, identifying the Dementia text in 75.3% and 70.7% of pairs, respectively (P = 0.44; Fig. 2e; Supplementary Table 4). A Bayesian equivalence analysis further supported comparable performance: the posterior probability that the true synthetic–real gap fell within a prespecified ±10-percentage-point margin was 0.85. Thus, synthetic narratives conveyed a clinically recognizable impairment signal comparable to that in real transcripts.

These results establish clinical alignment, generalizability, and specificity: the Dementia models reproduced real clinical linguistic patterns more closely than the controls, generalized across unseen discourse contexts, received impaired-range predicted MMSE scores, and remained clinically plausible to neurologists.

### DeLMs improve MCI classification through internal representations and synthetic data

We next asked whether DeLMs fine-tuned only on Pitt could generalize to a different population and a subtler level of cognitive impairment. To test this, we designed an MCI classification experiment using the Wisconsin Registry for Alzheimer’s Prevention (WRAP) cohort^30^, comprising 907 Cookie Theft transcripts – 787 from cognitively healthy individuals and 120 from individuals with MCI. This creates a challenging generalization setting: MCI is typically harder to detect from language than dementia^31^, and its prevalence in WRAP is only 13.2%, compared with the approximately balanced Pitt groups (Supplementary Table 1; Supplementary Note 4).

We used logistic regression classifiers under participant-grouped cross-validation, with held-out WRAP participants used only for testing. As WRAP-only baselines, we represent transcripts using either the 19 linguistic measures described above (*LING-19*) or pretrained sentence embeddings^32^, testing whether conventional linguistic or general-purpose language representations suffice without dementia-specific information. We also evaluated embeddings from the corresponding Vanilla checkpoints, to test whether general LLM representations alone accounted for any gain (Supplementary Table 5). We additionally evaluate random oversampling and SMOTE^33^ as class-imbalance controls that expand the MCI-class training set (Methods). These WRAP-only baselines performed near chance (ROC-AUC ≤ 0.56; PR–AUC ≤ 0.167, compared with a no-skill PR–AUC of 0.132).

We then designed three ways of incorporating information from the DeLMs – never trained on WRAP transcripts or any MCI data – into the classifiers (Fig. 3a). Geometry: rather than representing WRAP transcripts with linguistic measures or BERT-based embeddings, each WRAP transcript was passed through matched Healthy- and Dementia-tuned models, and the difference between their final-layer representations, averaged across tokens (Δh = h_Dementia_ − h_Healthy_), was used for classification. This tested whether Pitt-informed representations captured MCI-relevant information beyond WRAP-only representations. We report each family separately, alongside two combined variants: pooled prediction scores and concatenated representations before classification. A further variant added linguistic and perplexity features to test for complementary information (Fig. 3b; Methods). Augmentation: synthetic Cookie Theft descriptions generated by the Dementia models were added to the minority MCI class using the same LING-19 representation, either from each family separately or pooled across families. Fusion: Geometry and Augmentation prediction scores were averaged equally to test whether the two routes provided complementary information (Methods).

**Fig. 3.**
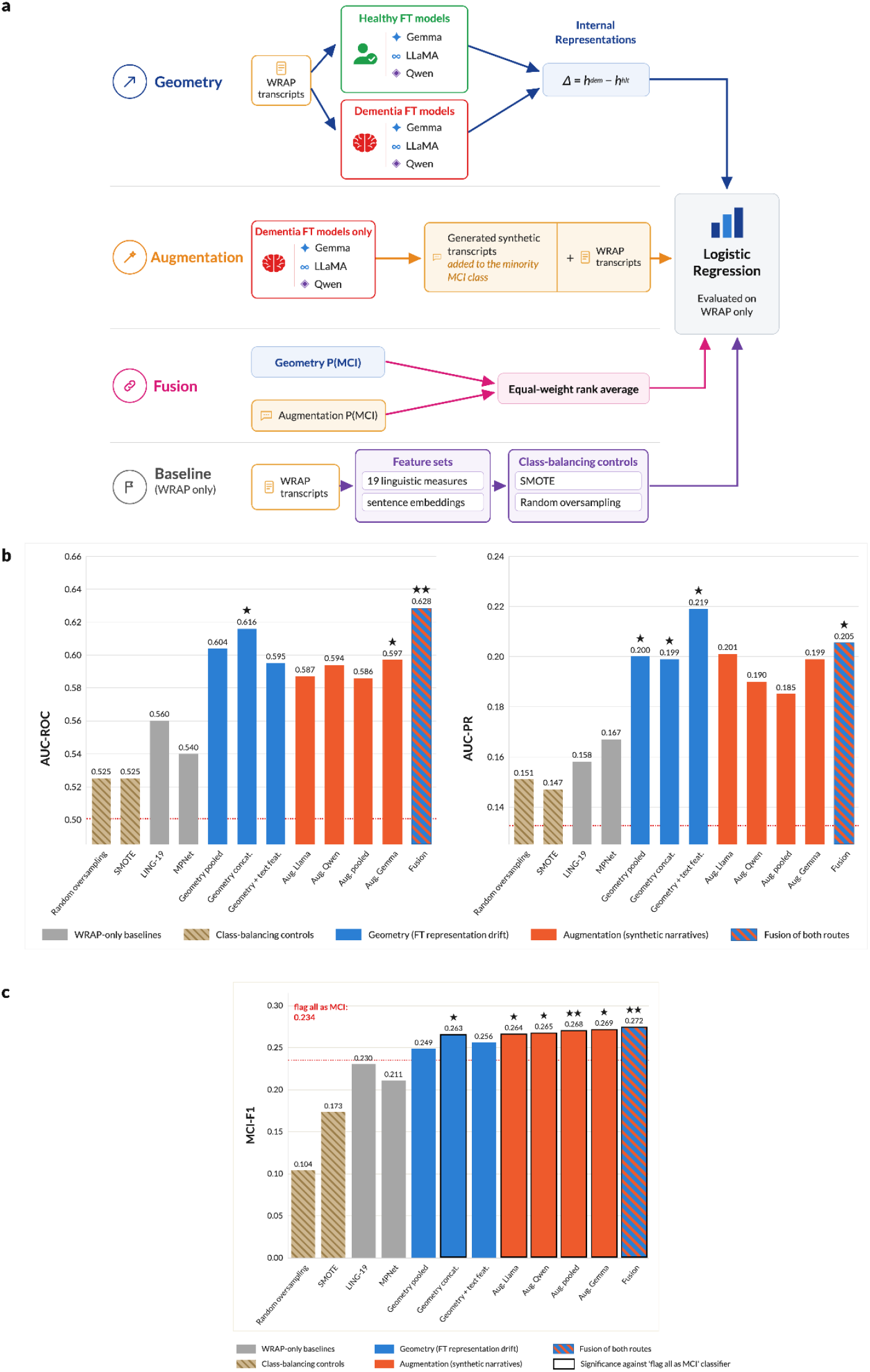
Generalization to the WRAP cohort. **a,** Three dementia-informed routes to MCI classification, compared with WRAP-only baselines and class-balancing controls. All methods use logistic regression trained on labeled WRAP data and evaluated by participant-grouped cross-validation; Pitt-trained language models remain frozen. Geometry: each transcript is passed through matched Healthy- and Dementia-tuned models, and the difference between their final-layer representations, averaged across transcript tokens (Δh = h_Dementia_ − h_Healthy_), is used for classification. Augmentation: synthetic Cookie Theft descriptions from Dementia-tuned models are added to the minority MCI class within training folds, using the same text representation as the baseline classifier. Fusion: Geometry and Augmentation scores are mapped to a common training-derived percentile scale and averaged equally. Baselines use 19 linguistic measures or pretrained sentence embeddings^32^; synthetic examples enter training data only. **b,** ROC-AUC and PR–AUC. Geometry variants: combining family-level predictions, concatenating representations, or adding text features. Pooled Augmentation uses texts from all three Dementia models. Random oversampling and SMOTE test whether rebalancing alone explains the gains. Stars: improvement over one (★) or both (★★) WRAP-only baselines. **c,** MCI-F1 at an 80%-sensitivity operating point. The red line marks the all-MCI reference (F1=0.234); stars indicate significant improvement over this reference.

The dementia-informed routes improved discrimination (Fig. 3b): fusion achieved ROC-AUC = 0.628 and PR–AUC = 0.205, while concatenated Geometry achieved ROC-AUC = 0.616 and PR–AUC = 0.199; the Geometry variant incorporating text features reached PR–AUC = 0.219. Augmentation provided modest performance in discrimination but improved minority-class detection: adding synthetic dementia narratives to the training data raised recall 2.2-fold relative to the same pipeline without augmentation (0.33 to 0.72). Random oversampling and SMOTE, matched to the same number of added training rows, did not reproduce this, indicating the gain reflects the added examples’ content rather than the shift in class ratio alone (Fig. 3b,c; Supplementary Table 5).

Together, these results show that our DeLMs encode information relevant to MCI classification in an independent cohort, both in their internal representations and their synthetic transcripts.

### Dementia models generalize from language to decision-making

To test generalizability beyond language, we asked whether DeLMs capture dementia-associated patterns in sequential decision-making, which is known to be affected by cognitive impairment^34–37^. For this experiment, we used the Zemla and Davis^38^ Iowa Gambling Task (IGT) dataset, which, among the studies we identified, was the only one providing complete trial-level trajectories for both MCI patients and healthy controls (45 per group). In the IGT, participants repeatedly choose among four decks with different reward and loss schedules, requiring them to learn from outcomes and adapt their choices^39^ (Supplementary Table 6). Participants respond with a single deck choice, without producing a verbal response. We subjected the Healthy- and Dementia-tuned models, as well as the controls, to the same game (Methods). Informed by established models of IGT behavior^40^, we represented games (real and synthetic) using 50 features summarizing deck preferences, sensitivity to reward/punishment, and sequential choice strategies (Fig. 4a; Methods; Supplementary Note 6). Fig. 4c illustrates two trial-level features from real and synthetic game sessions. For example, across all three model families, Dementia variants made fewer (or equivalent) value-maximizing choices than their Healthy counterparts, matching the direction observed in humans with MCI.

**Fig. 4.**
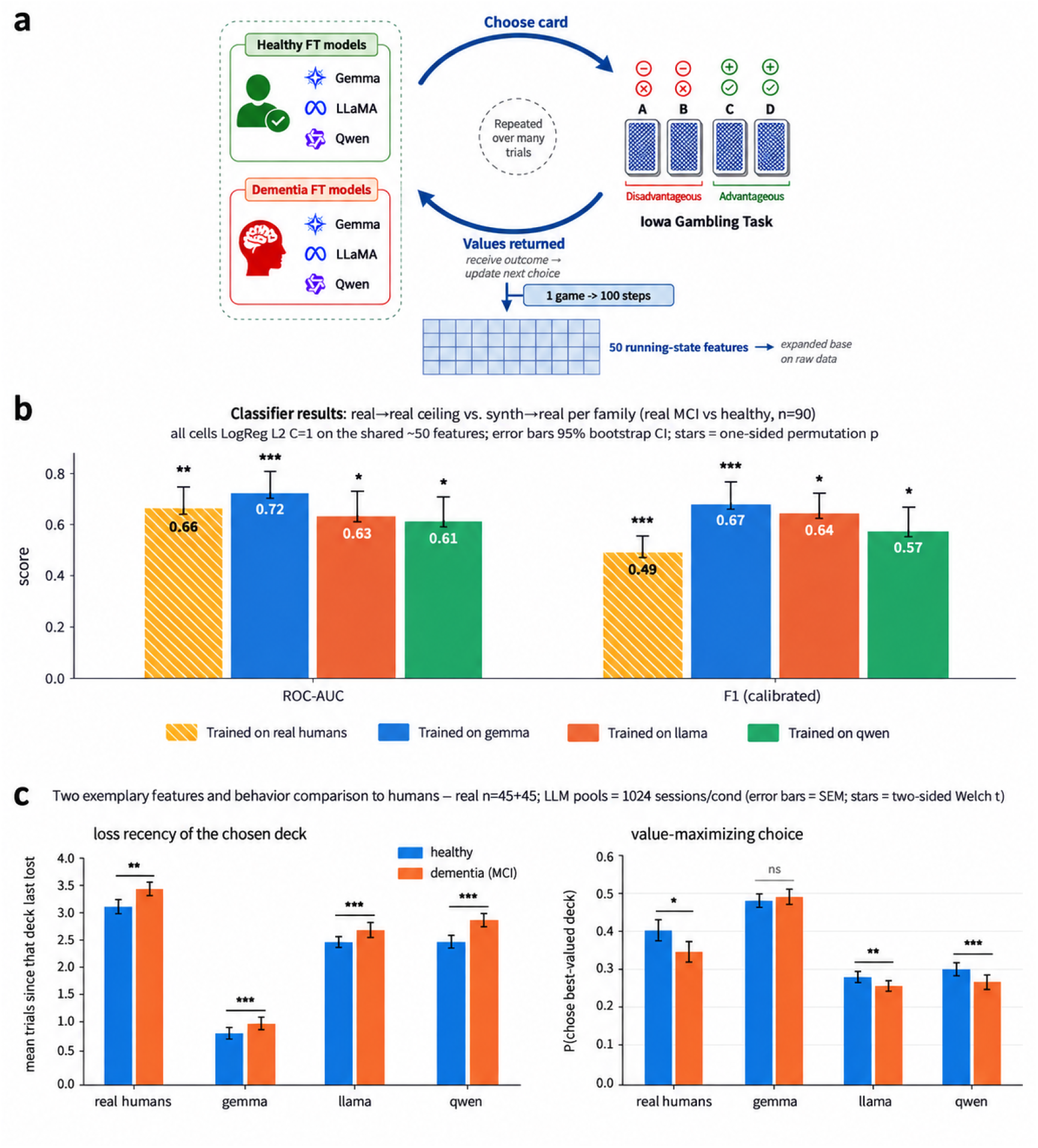
Dementia models show behavioral patterns that transfer from synthetic IGT play to real MCI. **a,** Experimental design. Healthy- and Dementia-tuned models completed repeated 100-trial Iowa Gambling Task sessions, choosing among four decks while receiving rewards, penalties, and updated game history. Sessions were represented using 50 behavioral features capturing choice and outcome dynamics. **b,** Synthetic-to-human transfer. Logistic-regression classifiers trained only to distinguish synthetic Healthy from Dementia sessions were applied unchanged to human sessions (45 healthy; 45 MCI). ROC-AUC and calibrated F1 are shown for Gemma, Llama, and Qwen, alongside a classifier trained directly on human data. Error bars show 95% bootstrap confidence intervals; stars denote one-sided permutation tests. **c,** Example behavioral features showing corresponding Healthy–Dementia/MCI differences in synthetic and real play: loss recency of the chosen deck and probability of choosing the currently highest-valued deck. Bars show means ± s.e.m. (real, n=45 per group; synthetic, n=1,024 sessions per condition); stars denote two-sided Welch t-tests (*P < 0.05, **P < 0.01, ***P < 0.001).

To evaluate our models’ performance in the IGT task, we followed Shapira et al.^41^ and assessed their predictive utility, asking whether their decision patterns can detect MCI in humans. For each model family, we trained a logistic-regression classifier to distinguish synthetic Healthy from Dementia game sessions and applied it directly, without exposure to human game sessions, to distinguish MCI from cognitively healthy participants (Methods). For comparison, a classifier trained directly on the human data achieves ROC-AUC = 0.657 under leave-one-out evaluation (P = 0.010), providing a human-trained benchmark (Methods).

Remarkably, classifiers trained only on DeLM decision-making distinguished human MCI from healthy participants across all three model families (ROC-AUC = 0.722, 0.629 and 0.611 for Gemma, Llama and Qwen; P = 0.0001, 0.0151 and 0.0348, respectively; Fig. 4b; Supplementary Table 7). Llama and Qwen were statistically indistinguishable from the human-trained benchmark, while Gemma numerically exceeded it, with overlapping confidence intervals. Neither control reproduced this pattern: classifiers trained on Healthy versus MediaSum or Healthy versus Magnitude-Matched performed at or below chance on the human cohort (ROC-AUC = 0.46–0.55 and 0.35–0.55, respectively).

### Dementia fine-tuning defines a structured, controllable axis in weight space

One might wonder whether DeLM behavior arises from degradation of the LLMs’ neural network itself – analogous to neural damage in dementia – rather than from learned adaptation. To examine whether this is true, we compared all fine-tuned models (Dementia, Healthy and MediaSum) with their Vanilla counterparts across three complementary properties: where weight changes occurred across network depth (weight-update magnitude across depth), how attention remained organized (attention entropy), and which layers contributed most strongly to model predictions (causal activation patching) (Methods). Across all three analyses, Dementia models closely resembled the other fine-tuned conditions: attention organization changed only minimally (normalized entropy differences ≤0.03; Fig. 5b; Extended Data Fig. 3b), the same layers contributed most strongly to predictions across conditions (r ≥ 0.998; Fig. 5a), and Healthy and Dementia models showed nearly identical depth-wise distributions of weight changes (r = 0.95–1.00; Methods; Extended Data Fig. 3a). Thus, DeLM behavior does not reflect broad computational or neural degradation. Instead, Healthy and Dementia tuning modified largely the same parts of an otherwise intact network, while moving their parameters to different values. This led us to ask whether the difference between the two models defines a meaningful direction in weight space.

**Fig. 5.**
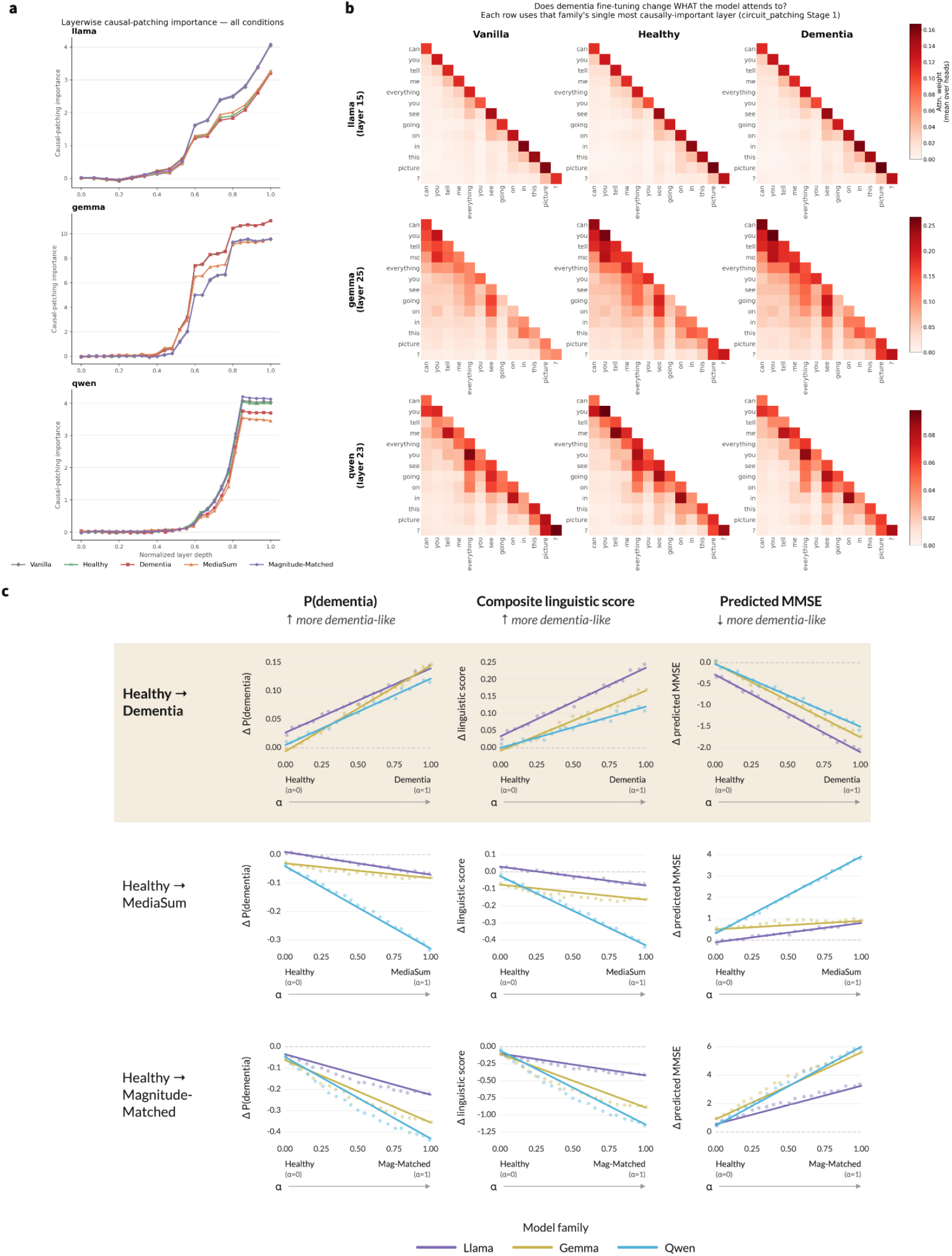
Preserved computational organization and a structured dementia-associated direction in weight space. **a,** Layerwise contribution to the target prediction across normalized network depth for Vanilla, Healthy, Dementia, MediaSum and Magnitude-Matched models, quantified using causal activation patching (Methods). Across model families, fine-tuning preserved the characteristic pattern of which layers contributed most strongly, with closely aligned profiles across conditions. **b,** Attention maps for a representative prompt at each family’s most influential layer, as identified by the causal-patching analysis in panel a. Vanilla, Healthy and Dementia models retained similar attention organization after fine-tuning; rows correspond to model families and columns to model conditions. **c,** Clinically grounded measures along linear weight-space trajectories from the Healthy checkpoint (α=0) toward the Dementia, MediaSum or Magnitude-Matched endpoint (α=1). Measures are shown as change relative to the corresponding Healthy endpoint and comprise p(dementia), a composite score summarizing 19 dementia-associated linguistic measures, and predicted MMSE. Points show mean values at each interpolation position and lines show ordinary least-squares fits; colors denote model families. Along the authentic Healthy→Dementia direction, all three measures changed approximately linearly toward a more dementia-associated profile, whereas both control directions changed smoothly but away from that profile.

For each LLM family, we created 19 intermediate models by taking a weighted average of the Healthy and Dementia weights, parameter by parameter. We denote the mixing proportion by α, where α = 0 is the Healthy model and α = 1 the Dementia model; for example, α = 0.1 corresponds to 90% Healthy + 10% Dementia weights. Including the Healthy and Dementia endpoints gave 21 evenly spaced models along this path. Each intermediate model generated 800 texts across the four narrative tasks. Evaluation spanned three measures: (i) a composite linguistic score summarizing the 19 measures into a single index of dementia-associated language relative to Healthy speech; (ii) p(dementia), from a classifier trained on real Healthy and Dementia transcripts; and (iii) predicted MMSE from the Pitt-trained ridge regressor. See Methods for full implementation details of the three measures.

All three measures changed approximately linearly along the continuum (R² = 0.816–0.975; Fig. 5c; Supplementary Table 8), in the expected direction: p(dementia) increased, linguistic behavior became more dementia-associated, and predicted MMSE decreased by 1.47–1.85 points. Thus, DeLM and Healthy models define a continuous, controllable axis of dementia-associated expression. Control interpolations (Healthy→MediaSum and Healthy→Magnitude-Matched) were also linear but moved away from the dementia-associated profile, indicating that linearity reflects smooth weight-space interpolation rather than the clinical signal itself^42^. Beyond α = 1, dementia-associated expression continued to increase until model breakdown by α = 4 (Extended Data Fig. 3d; Supplementary Note 8). Together, these findings identify a controllable dementia-associated direction in model weight space, providing a basis for studying how dementia-associated language varies continuously within a computational model.

## Discussion

In this study, we introduced Dementia Language Models: LLMs designed to learn a generalizable and controllable representation of cognitive impairment through language, together with an evaluation framework for establishing their validity and clinical grounding (Table 1). Across three distinct LLM families, fine-tuning on only a few hundred descriptions of a single picture yielded a coherent dementia-associated signal while preserving functional integrity. The signal showed clinical alignment across 19 linguistic measures, generalized to unseen narrative tasks, tracked predicted MMSE, and produced distinctions neurologists identified as readily as those in real transcripts. Generalizability extended beyond generated language and showed predictive utility in humans: DeLM internal representations improved MCI classification in an independent cohort, while synthetic decision-making patterns predicted MCI in real participants. Thus, although DeLMs operate through next-token prediction, the acquired signal was not confined to surface patterns of speech: it shaped both what the models encoded and how they behaved beyond language. Finally, the Healthy→Dementia weight difference defined a controllable continuum of dementia-associated expression, while neither control reproduced these effects, supporting their specificity to dementia-related learning.

**Table 1.** Mapping between the evaluation criteria and the experiments used to assess each criterion. Each criterion captures a distinct question about the DeLMs, with experiments spanning functional integrity, specificity, clinical alignment, generalizability, predictive utility and controllability.

| Evaluation criterion | Question | Experiments used to assess it |
| --- | --- | --- |
| **Functional integrity** | Did fine-tuning preserve general model function while inducing the intended dementia-associated signal without task-specific overfitting? | Instruction-following analysis; Cookie Theft leakage and memorization tests |
| **Specificity** | Are the observed effects specific to dementia-associated learning rather than generic fine-tuning, spoken-language adaptation, or parameter-change magnitude? | Comparison with MediaSum and Magnitude-Matched controls on all tasks |
| **Clinical alignment** | Do model differences correspond to patterns observed in real cognitive impairment? | Comparison with patient speech; MMSE predictions; neurologist discrimination |
| **Generalizability** | Does the learned signal extend beyond the training task and cohort? | Evaluation of generated narratives from unseen tasks (Cinderella retelling, sandwich preparation and general memories); evaluation in an independent MCI cohort (WRAP); non-linguistic decision-making (IGT) |
| **Predictive utility** | Can model-derived information help distinguish cognitive status in unseen individuals? | MCI classification in WRAP using model representations and synthetic-data augmentation; MCI prediction from synthetic IGT behavior |
| **Controllability** | Can the strength of the dementia-associated signal be varied systematically? | Healthy-to-Dementia weight-space interpolation and extrapolation, evaluated through language, predicted MMSE, and dementia probability |

Working closely with the models also made clear that signal consistency does not imply uniformity. Each family had its own sensitivities: Qwen responded more strongly to prompts matching the conversational register seen during fine-tuning, whereas Gemma was comparatively robust to such changes. Even in the IGT, eliciting deck choices by letter versus color could matter. These differences suggest that a consistent dementia-associated pattern can emerge through model-specific modes of expression. The signal also varied across narrative tasks (Fig. 2a). Because these tasks draw on different mixtures of memory, executive control, and narrative organization, such variation may ultimately help disentangle more specific dimensions of impairment rather than treating dementia-associated language as a single uniform signal.

Several limitations constrain our conclusions. The models were trained on a small English-language corpus from a single elicitation task and diagnostic process; behavioral generalization was tested on only one task; and MCI classification performance remained modest in absolute terms. Moreover, we do not claim that these LLMs have dementia, reproduce its biology or reveal disease mechanisms. They are computational models of patterns associated with patient language and behavior.

Finally, clinically convincing synthetic material carries ethical risks, including dataset contamination and fabricated patient records. Synthetic outputs should be clearly documented wherever they enter downstream datasets, and their use must respect the terms under which source data were collected. Any clinical application would require validation in the target population. Developed responsibly, Dementia Language Models could support controlled experiments, patient–clinician simulation and synthetic resources for underrepresented populations, providing an experimental complement to human research and reserving direct patient involvement for when it is truly required.

## Methods

### Models and fine-tuning

#### Base models

We used three open-weight, instruction-tuned language models from different families: Llama-3.2-1B-Instruct^16^ (meta-llama/Llama-3.2-1B-Instruct; 16 transformer layers, hidden size 2,048), Gemma-3-1B-IT^17^ (google/gemma-3-1b-it; 26 layers, hidden size 1,152), and Qwen2.5-1.5B-Instruct^18^ (Qwen/Qwen2.5-1.5B-Instruct; 28 layers, hidden size 1,536). Models in the 1–2B parameter range permitted full-parameter fine-tuning and direct weight-space analysis on the clinical corpus.

Instruction-tuned models were used because their chat-based format supports controlled elicitation at evaluation time. Using three distinct model families allowed us to test whether the observed effects generalized beyond a single architecture. The original checkpoint of each family is referred to as the Vanilla model.

#### Pitt corpus

Clinical fine-tuning data were drawn from the Pitt corpus of DementiaBank^15^, a controlled-access repository available to researchers under the TalkBank Code of Ethics. The Healthy set comprised 232 transcripts from 99 participants with a Control diagnosis, and the Dementia set comprised 217 transcripts from 141 participants with probable Alzheimer’s disease. Mean transcript lengths were 99.7 and 94.4 words, respectively. Because some participants contributed repeated visits, transcript counts exceed participant counts; each transcript was treated as one fine-tuning example. Ages at the corresponding visits were 64.8 ± 7.8 years for Healthy and 71.6 ± 8.5 years for Dementia. Transcript-level observations comprised 146 female and 86 male observations in Healthy and 143 female and 74 male observations in Dementia. Because age and sex could contribute to linguistic differences independently of diagnosis, the principal linguistic analyses were repeated with these variables included as covariates (Supplementary Table 1; Supplementary Note 1; Supplementary Table 2).

Throughout the study, Dementia refers to participants labelled Probable Alzheimer’s disease in Pitt, based on the research-grade NINCDS–ADRDA diagnostic criteria^43^: clinical examination and mental-status assessment, supported by neuropsychological evidence of impairment in at least two cognitive domains, progressive worsening of memory and other cognitive functions, absence of disturbed consciousness, and no alternative systemic or neurological disorder sufficient to explain the deficits.

#### Transcript preprocessing and formatting

CHAT transcription markup was removed or normalized before fine-tuning. Verbalized hesitations and unintelligible-speech placeholders were retained, unfilled-pause annotations were converted to orthographic ellipses, and interviewer backchannel markers were stripped; full preprocessing details are provided in Supplementary Note 2.

Each transcript was formatted as a single-turn user–assistant exchange using the model’s native chat template. For the Pitt corpus, the user turn contained one of three fixed elicitation prompts, sampled independently at random for each training example to reduce sensitivity to the exact wording of the prompt: “Describe the cookie theft picture.” / “Please describe what you see in detail.” / “Describe the picture.” The assistant turn contained the corresponding participant transcript.

No explicit system prompts were supplied. Inputs were formatted using each model’s native chat format, which may add default system content; these automatically added tokens were excluded from the training loss. Hence, differences in input formatting could not account for within-family system prompt contrasts.

#### Fine-tuning procedure

For each model family, we created a Healthy and a Dementia variant by full-parameter fine-tuning on the corresponding Pitt transcript set. All model parameters were updated; no adapters or other parameter-efficient fine-tuning methods were used. Training loss was computed only over assistant-response tokens. Labels corresponding to the user turn and any template-injected system tokens were masked. The end-of-turn token was included in the supervised span so that the models learned to terminate their responses.

Models were optimized with AdamW (β_1_ = 0.9, β_2_ = 0.999), a learning rate of 1 × 10^−5^, and gradient clipping at 1.0 for up to six epochs. The batch size was 2. Effective batch size was 16 for Llama and 4 for Gemma and Qwen, achieved through gradient accumulation. Validation loss was evaluated every 50 optimizer steps and checkpoints were saved at the same interval. For each run, the checkpoint with the lowest validation loss was retained. Remaining hyperparameters, including weight decay, warmup ratio, and learning-rate schedule, were selected separately for each model family through search according to validation loss.

Fine-tuning was performed using the Hugging Face Transformers Trainer (v4.55) with PyTorch 2.9 on a single NVIDIA RTX A6000 GPU (48 GB; Ampere architecture). Training was performed end-to-end in bfloat16 precision: model weights, gradients, and AdamW moment estimates were stored in bfloat16, with no fp32 master weights and no mixed-precision autocast wrapper. Gradient checkpointing was not required. The same precision and optimization regime was applied to all fine-tuned conditions, including the control models.

#### Data splitting and multiseed averaging

For each fine-tuning run, transcripts were divided into 80% training and 20% validation sets. Splits were constructed such that all transcripts from the same participant remained within a single partition, preventing repeated visits from the same individual from appearing in both training and validation. A new split was generated independently for each random seed.

Each fine-tuning condition was trained independently using three random seeds (42, 123, and 7), each with its own train–validation split and early stopping. The best-validation-loss checkpoints from the three runs were then merged by element-wise averaging of corresponding parameter tensors, following established weight-averaging practice to negate seed sensitivity^42^. Because all runs within a condition originate from the same pretrained initialization, corresponding parameters are directly aligned for averaging. All reported analyses therefore use the resulting multiseed-averaged models.

#### Control models

We constructed two control conditions to distinguish dementia-associated learning from more general consequences of fine-tuning.

Magnitude-Matched control: For each model family, we computed the parameter update produced by Dementia fine-tuning:

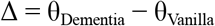

where θ_Dementia_ is the multiseed-averaged Dementia checkpoint.

For each parameter tensor independently, the corresponding delta was converted to float32, flattened, randomly permuted, reshaped to its original dimensions, and added back to the Vanilla parameters:

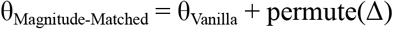

A new permutation was drawn independently for each tensor using a fixed random seed (seed 1), yielding one Magnitude-Matched control model per family.

Permutation preserves the exact set of update values within each tensor and therefore its norm, mean, variance, and empirical distribution, while destroying the organization of those values across parameters. Because permutation is performed separately within each tensor, the amount of update assigned to each layer and submodule is also preserved. The control therefore matches both the magnitude of Dementia fine-tuning and its tensor- and depth-wise allocation through the network while removing its learned parameter-level organization. The Magnitude-Matched condition should therefore be interpreted as a matched-size, matched-allocation unstructured perturbation control.

MediaSum control: This control used 500 spoken interview responses randomly selected from MediaSum, a corpus of NPR and CNN interview transcripts^20^. This control tests whether effects attributed to clinical fine-tuning could instead be explained by adaptation to spontaneous spoken language or interviewer–respondent discourse more generally. Each Vanilla model was fine-tuned on this corpus using the same overall procedure as the Pitt variants, including full-parameter fine-tuning, independent train–validation splits and early stopping for each of three random seeds, and element-wise checkpoint averaging. Hyperparameters were selected using the same validation-loss-based procedure as for Pitt.

MediaSum transcripts were processed to approximate the instruction–response structure of the Pitt examples. Interviewee turns from non-broadcaster speakers were extracted as candidate assistant responses, while broadcaster roles were excluded using speaker-role pattern matching. Responses shorter than 20 words were discarded. The immediately preceding interviewer turn was retained as the user message when it consisted of a 4–10-word question ending in a question mark. To reduce over-representation of individual speakers, no more than three exchanges were retained from any one speaker. Each resulting training example therefore paired a real interviewer question with the corresponding interviewee response.

The same 500 MediaSum exchanges were used for all three model families. For each fine-tuning seed, exchanges were independently divided into 80% training and 20% validation sets at the exchange level. Speaker identity was not retained in the processed corpus and therefore could not be used to group the train–validation split.

The MediaSum examples approximated the Pitt data in elicitation structure, response length, and training-set size. Retained MediaSum responses averaged 82.0 words (median 78), compared with 99.7 words for Healthy and 94.4 words for Dementia transcripts. Whereas Healthy and Dementia speech came from the same clinical corpus and elicitation protocol, MediaSum provided a non-clinical control for adaptation to spontaneous, interviewer-elicited speech. Together with the Magnitude-Matched control, this allowed us to distinguish dementia-associated learning from effects attributable to spoken-language register, elicitation format, or fine-tuning more generally.

### Functional integrity evaluation

We evaluated the effects of fine-tuning on general instruction-following ability, acquisition of Pitt speech characteristics, and Cookie Theft-specific leakage.

#### Evaluation set

We evaluated every model variant on the same in-house set of 300 prompts designed to assess general instruction-following ability. The evaluation framework was informed in part by the types of instruction constraints represented in IFEval^44^, including length, keyword, structural, and output-format requirements, and was extended with prompts assessing general knowledge, reasoning, and robustness to typographical errors.

The prompts covered seven categories: general-knowledge questions (n = 45); length constraints, such as exact, minimum, or maximum word or sentence counts (n = 45); keyword constraints, requiring or prohibiting particular terms (n = 30); structural constraints, such as prescribed opening or closing text, list structure, or multipart responses (n = 42); output-format constraints, including JSON or CSV formatting, capitalization, and exact-copy instructions (n = 52); reasoning tasks, including arithmetic, parsing, and multistep composition (n = 58); and robustness probes, including tolerance to typographical errors (n = 28). The complete evaluation set is provided in Supplementary Data 1.

Each of the 15 model variants answered all 300 prompts, yielding 4,500 responses. Sampling was disabled, and generation used a maximum of 128 new tokens and a repetition penalty of 1.1, applied identically across models and conditions.

#### Evaluation criteria

Each response was evaluated independently along three axes: instruction adherence, spontaneous-speech markers, and Cookie Theft-specific bleed. The three criteria were scored in separate judge calls.

Instruction adherence measures whether the response understood and satisfied the requested task and was scored on a five-point scale. Reference answers and task-specific constraints associated with each prompt were provided to the judge as reference information for assessing correctness and compliance. A graded scale was used rather than exact reference-answer matching alone to distinguish complete adherence from partial compliance, minor technical violations, substantive errors, and fully nonresponsive outputs. The rubric was: 5, fully satisfies the requested content and all constraints; 4, satisfies the task with only a trivial technical error; 3, provides substantively correct content but violates a hard constraint, such as an incorrect sentence count or missing required keyword; 2, engages with the requested task but gives substantively incorrect content; and 1, produces an irrelevant, incoherent, or otherwise nonresponsive answer.

Conversational fillers and other stylistic characteristics acquired during fine-tuning were not penalized unless they caused the response to violate the instruction.

Spontaneous-speech markers were scored as a binary indicator of whether the response exhibited more than one characteristic associated with the speech register represented in the Pitt corpus. The annotation rubric included conversational fillers and hesitation markers, discourse markers, spoken-language phrasing, perseverative or unnecessary repetition, off-task conversational material, low-information or empty speech.

Cookie Theft bleed was evaluated separately as a more specific measure of fine-tuning-task leakage. A response was flagged when it introduced content from the Cookie Theft picture into an unrelated response, for example by describing a boy taking cookies from a jar while standing on a stool or a sink overflowing while a woman washes dishes. Incidental use of overlapping vocabulary in an appropriate context – for example, answering “cookies” when asked to name a sweet food – was not considered bleed.

#### Validation of LLM-based annotation

Scoring the complete evaluation set required 15 model variants × 300 responses × 3 criteria = 13,500 annotations. We therefore used an LLM judge for full-scale annotation and validated its performance against human annotators.

Four human annotators independently evaluated the same subset of 50 model responses using the same annotation rubrics. Responses were presented blinded to model identity and condition and in randomized order. Inter-annotator agreement was Fleiss’ κ = 0.881 for instruction adherence, κ = 0.749 for spontaneous-speech markers, and κ = 0.80 for Cookie Theft bleed.

We compared candidate LLM judges using the Alternative Annotator Test^22^. For each human annotator in turn, that annotator was held out, and the held-out human and candidate LLM judge were independently compared against the remaining human annotators. Agreement was quantified using negative root-mean-square error for the 1–5 adherence ratings and exact-match accuracy for the two binary criteria. Non-inferiority was evaluated with a disagreement tolerance of ε = 0.15 and FDR correction at q = 0.05. For each criterion independently, a winning rate of at least 0.5, corresponding to successful statistical replacement of at least half of the human annotators, was used as the criterion for selecting an automated judge.

GPT-5.4-mini (gpt-5.4-mini-2026-03-17)^21^ satisfied the selection criterion on all three measures and was therefore used to annotate the complete evaluation set. Full-scale annotation used temperature 0 and structured outputs, with each criterion evaluated in a separate call.

#### Statistical analysis of functional integrity

Analyses were performed separately within each model family, using the corresponding Vanilla model as the reference condition. Because all model variants answered the same 300 prompts, comparisons were paired by prompt. Instruction-adherence scores were compared with Vanilla using paired t-tests. Binary spontaneous-speech-marker and Cookie Theft-bleed outcomes were compared using exact McNemar tests (Extended Data Fig. 1).

Effect sizes for instruction adherence were reported as paired Cohen’s d with 95% confidence intervals, and effect sizes for the binary criteria as risk differences (RD) with 95% confidence intervals. Because Cookie Theft bleed was rare, raw counts and proportions are also reported alongside statistical comparisons. All statistical tests were two-sided.

### Narrative generation and linguistic analysis

#### Narrative generation

We elicited four narrative types spanning discourse tasks used in clinical language assessment: Cookie Theft picture descriptions (scene description; the fine-tuning task), Cinderella story retellings (narrative recall)^45^, sandwich-preparation instructions (procedural discourse)^45^, and General Memories (open-ended autobiographical narration)^46^. For each of the 15 model variants and each task, we generated 200 responses using a bank of 10 manually written prompt formulations per task (Supplementary Data 2). The formulations requested the same underlying discourse while varying the wording of the instruction, to guard against prompt sensitivity.

Prompts were presented as a single user turn with no system prompt. Temperature was sampled independently for each generation from {0.6, 0.8, 0.9, 1.0, 1.1}; nucleus sampling used top-p = 0.95, with a maximum of 200 new tokens. No repetition penalty was applied.

To guard against memorization, every Cookie Theft narrative generated by the Healthy- and Dementia-tuned variants was compared with the corresponding Pitt fine-tuning transcripts. Responses sharing any exact contiguous 10-word sequence with a training transcript were discarded and regenerated. Pitt utterances have a median length of six words and a 75th percentile of nine words, making a 10-word match unlikely to arise from short phrases naturally shared across descriptions of the same picture (e.g., “mother is washing dishes”). No narrative met this memorization criterion (Supplementary Note 3).

#### Linguistic feature extraction

Generated narratives were represented using 19 linguistic measures previously associated with dementia-related language, spanning four domains. All literature sources, measure directions, definitions and formulas can be found in Supplementary Table 2. Lexical diversity and richness included moving-average type-token ratio (MATTR; window = 20), Brunet’s index, Honoré’s statistic, and unigram and bigram repetition intervals. Syntactic and informational complexity included Flesch-Kincaid grade level, propositional idea density, and mean word length. Lexico-grammatical and referential specificity included noun frequency, verb frequency, noun-to-verb ratio, pronoun-to-noun ratio, adposition frequency, and mean lexical frequency across all words, content words, nouns, and verbs.

Disfluency was represented by filled-pause rate and empty-pause rate, the latter defined from orthographic ellipses in the original text.

Text processing used spaCy with the en_core_web_sm pipeline for tokenization, part-of-speech tagging, and sentence segmentation; lexical-frequency estimates were obtained using wordfreq, which incorporates SUBTLEX frequency information alongside other corpus sources; and syllable counts were approximated using pyphen.

Measures undefined for a particular narrative, such as repetition intervals when no item recurred or ratio measures with an empty denominator, were retained as missing during feature extraction (Supplementary Note 9). Downstream analyses requiring complete numeric input handled missing values as described for the corresponding analysis.

Before examining model outputs, each measure was assigned a prespecified Healthy-to-Dementia direction based on previous clinical and computational literature (Supplementary Table 2). For example, a higher pronoun-to-noun ratio was prespecified as dementia-associated based on previous findings of reduced referential specificity in Alzheimer’s disease^47^.

Complete metric definitions, implementation details, direction assignments and literature grounding are provided in Supplementary Table 2.

#### Directional consistency across narrative tasks

We first tested whether the linguistic contrast induced by Healthy versus Dementia fine-tuning followed the clinically expected direction and generalized beyond the Cookie Theft fine-tuning task. Directionality was evaluated separately for each model family and narrative task, yielding 12 family-by-task combinations (three families × four tasks).

Within each combination, we compared the mean of each of the 19 measures between Healthy- and Dementia-tuned generations. A measure was counted as directionally consistent when the sign of the Healthy minus Dementia difference matched its prespecified clinical direction. The number of consistent measures out of 19 was compared with the chance expectation of 0.5 using a one-sided exact binomial test. Because the linguistic measures are correlated, these tests were treated as summary tests of directional consistency; quantitative fidelity to the human contrast was assessed separately below. A correlation matrix among the 19 features can be found in Extended Data Fig. 2.

The same procedure was applied to the Healthy-MediaSum and Healthy-Magnitude-Matched contrasts to test whether similar directional shifts arose from conversational fine-tuning or parameter change alone.

#### Clinical fidelity to real Pitt speech

Quantitative fidelity analyses were restricted to Cookie Theft, the only task with directly matched real patient speech. Because model families differ in their baseline linguistic profiles, analyses focused on the within-family Healthy-Dementia contrast rather than on absolute synthetic-human differences. We assessed the magnitude of individual linguistic effects, equivalence of the full contrast, and alignment of the 19-dimensional linguistic profile.

#### Difference-of-differences

For each model family and linguistic measure, we calculated the synthetic Healthy-Dementia difference and compared it with the corresponding Real Healthy-Real Dementia difference. The difference between these two contrasts is zero when the synthetic effect exactly matches the real clinical effect. Uncertainty was estimated using 5,000 non-parametric bootstrap resamples. Two-sided p-values were corrected across the 19 measures within each model family using the Benjamini-Hochberg FDR procedure. A non-significant difference was interpreted as an absence of detectable mismatch, not as evidence of equivalence. The same analysis was applied to the Healthy-MediaSum and Healthy-Magnitude-Matched contrasts.

#### Joint Bayesian relative-fidelity analysis

We additionally quantified whether the multivariate mismatch between the synthetic and real Healthy–Dementia contrasts was smaller than the magnitude of the real clinical contrast itself. Mismatch across the 19 direction-aligned measures was aggregated into a composite error and compared with δ, the magnitude of the real Healthy–Dementia contrast. An error/δ ratio of 1 therefore represents a mismatch as large as the real clinical effect, whereas values below 1 indicate progressively closer agreement. Because δ was estimated from the same Pitt sample, uncertainty in both quantities was propagated using a Bayesian bootstrap with 8,000 posterior draws, recomputing the error and δ in each draw and reporting P(error < δ). Repeated Pitt visits were collapsed to participant-level means, and the same analysis was applied to the Vanilla, MediaSum, and Magnitude-Matched controls.

#### Contrast-vector cosine similarity

We quantified alignment between the synthetic and real Healthy-Dementia contrast vectors using cosine similarity. Before constructing the vectors, each measure was scaled by its pooled within-group standard deviation across Real Healthy, Real Dementia, Synthetic Healthy, and Synthetic Dementia observations. Unlike the equivalence analysis, which evaluates the magnitude of the mismatch, cosine similarity captures the relative pattern of effects across measures independently of overall scale. Confidence intervals were estimated from 10,000 stratified bootstrap replicates, resampling narratives within each group and recomputing the scaling factors and contrast vectors in every replicate. We report percentile 95% confidence intervals, with intervals excluding zero indicating positive alignment. The same procedure was applied to the control contrasts.

#### MMSE prediction from linguistic measures

To test whether the linguistic changes induced by dementia fine-tuning mapped onto a clinically interpretable measure of cognitive severity, we trained a Ridge regressor to predict Mini-Mental State Examination (MMSE)^29^ scores from the same 19 linguistic measures. The analysis included 399 Pitt Cookie Theft transcripts from 241 participants diagnosed as “control” or “ProbableAD” with a non-missing MMSE score recorded at the same visit (185 Control, 214 ProbableAD). Multiple visits were retained, but all visits from a participant were assigned to the same cross-validation fold. The regression pipeline used median imputation and feature standardization fitted within each training fold; the regularization parameter was selected from 13 logarithmically spaced values between 0.01 and 10,000 using inner three-fold participant-grouped cross-validation, minimizing RMSE.

Generalization performance was evaluated using outer five-fold participant-grouped cross-validation, stratified by the MMSE severity band of each participant’s first available visit: 0–17, moderate-severe impairment; 18–23, mild impairment; and 24–30, no impairment.

Predictions were clipped to the valid MMSE range of 0–30. Using the prespecified fold seed of 42, out-of-fold performance was RMSE 5.45 (95% CI, 5.06–5.84), MAE 4.46, R² 0.30, Spearman ρ = 0.56, and quadratic-weighted κ = 0.47. This was of a similar order to prior work predicting MMSE from Pitt or related Cookie Theft transcripts^48,49^ providing a validity check on the scale of prediction error rather than a direct benchmark. Confidence intervals were estimated using 1,000 participant-cluster bootstrap samples, and repeating the complete procedure across five fold seeds (42–46) yielded stable performance (RMSE 5.42 ± 0.04; Supplementary Table 3).

We then refit a final Ridge model on all 399 eligible Pitt transcripts using the same preprocessing pipeline, with the regularization parameter re-selected by participant-grouped cross-validation on the full cohort. This fitted model was then fixed and applied unchanged to all synthetic narratives. For real Pitt reference scores, we used only out-of-fold predictions, ensuring that no transcript was scored by a model trained on that participant. Synthetic results were summarized separately by model family and pooled across Llama, Gemma, and Qwen, with each family contributing 200 generations per condition and task.

Predicted MMSE <24 was defined as the primary impaired-range threshold^29^, with progressively stricter thresholds of <22 and <20 used to test whether Healthy-Dementia separation increased with predicted severity. For each threshold, we calculated the relative risk of falling below the cutoff for Dementia- versus Healthy-tuned generations; confidence intervals used the Wald approximation on the log scale, and significance was assessed with Fisher’s exact test on the corresponding 2 × 2 contingency table.

#### Neurologist discrimination study

Five expert neurologists, all co-authors, independently completed a two-alternative forced-choice task comprising 30 synthetic and 30 real Healthy–Dementia pairs. At the time of evaluation, the neurologists had not participated in study design or model development and were blinded to all information about model training, model identity, prompting, preprocessing, generation procedures, and the broader study beyond the instructions required to complete the task. They were shown only the paired texts and became study co-authors only after their ratings had been collected. Synthetic pairs contained one Healthy- and one Dementia-generated narrative from the same model family and elicitation task; real pairs contained transcripts from one cognitively healthy and one cognitively impaired speaker in the same elicitation setting. Within each pair, texts were matched on word count and disfluency rate by minimizing a normalized word-count and filler-count difference score and were screened for task adherence. For each neurologist, pair order and the left–right position of the impaired text were independently randomized, and Healthy/Dementia labels were hidden. Raters were informed that spoken-language features such as hesitations, disfluencies and informal phrasing could occur in both texts and were asked to select the narrative showing greater evidence of cognitive impairment (Supplementary Table 4).

Accuracy was calculated separately for synthetic and real pairs and compared using Fisher’s exact test on pooled correct and incorrect judgments. As a complementary analysis, we quantified uncertainty in the difference in detection accuracy (synthetic minus real) using independent beta-binomial models for the two conditions. We evaluated the posterior probability that the absolute difference fell within ±10 percentage points and, as a stricter sensitivity analysis, ±7 percentage points. Posterior estimates were based on 400,000 Monte Carlo draws and evaluated under both uniform Beta(1,1) and Jeffreys Beta(0.5,0.5) priors.

#### Statistical reporting of linguistic analysis

All tests were two-sided unless a directional hypothesis was prespecified. Statistical significance was evaluated at α = .05. Where the 19 linguistic measures were tested simultaneously, p-values were adjusted within each model family using the Benjamini-Hochberg false-discovery-rate procedure. Bootstrap confidence intervals were percentile intervals unless otherwise specified, and effect sizes and confidence intervals were reported alongside significance tests.

### MCI classification in the WRAP cohort

We tested whether dementia-associated information learned from Pitt could support classification of MCI versus Healthy in an independent cohort. Two complementary routes used the fixed fine-tuned models: Geometry extracted information from their internal representations, whereas Augmentation used their generated language. Fusion combined the resulting predictions. The language models were trained exclusively on Pitt and were never fine-tuned on WRAP transcripts or MCI labels; WRAP data was used to train and evaluate only the downstream classifiers.

#### Cohort and preprocessing

We used Cookie Theft picture descriptions from the Wisconsin Registry for Alzheimer’s Prevention (WRAP)^30^, an independent longitudinal cohort enriched for Alzheimer’s disease risk. The analysis included the latest available transcript from each of 907 participants: 787 cognitively healthy individuals and 120 with MCI by consensus clinical diagnosis (13.2% prevalence; Healthy:MCI ratio of 6.6:1). Participants with a dementia diagnosis (n = 35) were excluded from this Healthy-versus-MCI analysis (Supplementary Note 4).

Raw CHAT transcripts were processed using the same conventions as the Pitt data: markup and annotation codes were removed, while hesitations, retraces and repetitions were retained. No WRAP transcript was used to fine-tune the language models or to generate synthetic narratives.

#### Evaluation design

All downstream methods were evaluated using the same outer five-fold cross-validation split, stratified by cognitive-status label and grouped by participant (seed 42). Each participant therefore contributed one held-out prediction per method, and comparisons were paired across the same 907 participants. Hyperparameters were selected within each outer training fold using a grouped three-fold grid search, with average precision as the selection criterion. Median imputation, standardization and PCA, where applicable, were fitted using training data only.

#### WRAP-only baselines

Two WRAP-only baselines represented transcripts using either (i) the 19 linguistic measures described above (LING-19) or (ii) frozen pretrained sentence embeddings from MPNet (all-mpnet-base-v2^32^), a BERT-based sentence-embedding model. Additional experiments were conducted using RoBERTa-base^50^, LLM2Vec-Llama^51^, and embeddings from the Vanilla checkpoints (Llama-3.2-1B-Instruct, Gemma-3-1B-IT, and Qwen2.5-1.5B-Instruct). The best-performing, and therefore hardest-to-beat, baseline was MPNet. WRAP baselines used logistic regression with balanced class weights, with regularization strength selected within the inner grid search. All preprocessing and classifier fitting were performed exclusively on WRAP training participants within the cross-validation procedure.

#### Geometry: classification from internal representation shifts

Geometry tested whether the internal representations of the fine-tuned models contained information relevant to MCI classification. Each WRAP transcript was passed through the matched Healthy- and Dementia-tuned checkpoints of a model family, using the Cookie Theft task prompt and the model’s native chat format. We extracted a vector representation of the transcript from the final hidden layer by averaging across its transcript tokens, excluding prompt tokens from pooling. Representations were stored in float32.

For each transcript, the classification feature was the difference between the two representations:

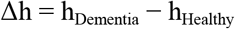

The difference vectors contained 2,048, 1,152 and 1,536 dimensions for Llama, Gemma and Qwen, respectively. The primary Geometry configuration concatenated the three family-specific vectors into a 4,736-dimensional representation, followed by within-fold PCA and logistic regression with balanced class weights. The number of PCA components was selected from {8, 16, 32, 64}, subject to a cap of 0.6 × the number of training observations, and logistic-regression regularization strength was selected within the inner grid search.

We also evaluated individual model families, score-level pooling and an extended configuration incorporating linguistic and perplexity features. Score-level pooling averaged the three families’ predicted probabilities, rather than averaging their raw representation vectors. No synthetic narratives were used in Geometry, and the language-model checkpoints remained fixed throughout.

#### Augmentation: classification using generated language

Augmentation tested whether language generated by the Pitt-trained Dementia models could improve minority-class learning. Synthetic narratives were represented using the same 19 linguistic measures (LING-19) as the corresponding WRAP-only baseline and added to the minority MCI class within each outer training fold. Classification used logistic regression with balanced class weights, with regularization strength selected within the inner grid search.

Augmentation used only features that could be computed comparably for synthetic and real WRAP transcripts. Internal model representations and model-derived perplexity features were therefore excluded from this route. Using the same checkpoint to generate a synthetic narrative and subsequently represent or score it would introduce model-specific self-generation information that is unavailable for the real WRAP transcripts being classified, confounding the intended comparison between synthetic and human language. These features were instead evaluated in the Geometry route.

The three Dementia-tuned model families generated 200 Cookie Theft narratives each. The final pooled augmentation analysis used 600 synthetic narratives, which were added to the MCI class of each outer training fold. Because the generators were trained on a Healthy–Dementia contrast, these examples were used as a proxy for the cognitive-impairment direction rather than as simulations of MCI itself. Whether this information could improve the subtler Healthy-versus-MCI classification was part of the generalization test.

To distinguish dementia-generated information from generic rebalancing, we evaluated balanced class weighting, random oversampling and SMOTE^33^, as well as augmentation using narratives generated by the Vanilla and Healthy-tuned models. All comparisons used the same LING-19 representation and logistic-regression pipeline. Oversampling, SMOTE and Dementia augmentation added the same fixed number of rows within each corresponding training fold (600 for the pooled source). A supplementary analysis additionally pooled dementia-generated narratives from all four elicitation tasks rather than Cookie Theft alone (Supplementary Note 5).

Synthetic augmentation, random oversampling, and SMOTE were performed independently within each inner-training split during hyperparameter selection; no synthetic or resampled observations derived from an inner-validation participant entered the corresponding inner-training set. After hyperparameter selection, the selected procedure was reapplied to the complete outer-training fold, the classifier was refit, and performance was evaluated on untouched real participants in the outer test fold. Synthetic examples never entered outer test folds.

#### Fusion of Geometry and Augmentation

Fusion tested whether the representation-based and generation-based routes provided complementary predictive information. The combination paired the selected concatenated Geometry configuration with pooled Dementia augmentation. The equal-weight combination rule and pairing were specified before fusion comparisons.

Within each outer fold, prediction scores from Geometry and Augmentation were mapped to percentile ranks using empirical score distributions fitted on the corresponding training partition. These training-derived mappings were then applied to held-out scores, and the two percentile values were averaged with equal weights. No additional classifier or outcome-dependent fusion weight was fitted.

### Evaluation metrics and operating-point selection

Ranking performance was assessed using ROC-AUC and PR–AUC (average precision). PR–AUC complements ROC-AUC by emphasizing performance on the minority MCI class and by providing a prevalence-dependent reference under class imbalance. The no-skill references were 0.5 for ROC-AUC and 0.132 for PR–AUC, corresponding to the MCI prevalence.

We additionally reported MCI-class F1, precision, recall and balanced accuracy at a screening-oriented operating point targeting 80% sensitivity. Within each outer fold, the final fitted classifier produced scores for the real WRAP training participants. The threshold was selected as the highest observed score threshold achieving at least 80% sensitivity among training positives: positive-class scores were sorted in ascending order, the index was set to floor(0.2 × n_positive_), and the corresponding score was used as the threshold. Participants with scores greater than or equal to this threshold were classified as MCI; ties were therefore included. The selected thresholds were applied unchanged to held-out participants, and realized test sensitivity was allowed to differ from the 80% training target.

MCI-F1 was used as the primary F1 measure because MCI was the target class. For descriptive context, a classifier labelling every participant MCI would attain precision equal to the cohort prevalence, recall of 1 and MCI-F1 of 0.234.

#### Statistical analysis of WRAP generalization

For each method, 95% confidence intervals were estimated using class-stratified participant bootstrap with 1,000 resamples. Pairwise method comparisons used 50,000 paired participant-bootstrap resamples. Within each comparison, the same resampled participant indices were applied to both methods’ held-out predictions, preserving their pairing. We report 95% percentile intervals for between-method differences and two-sided, sign-based P values. Primary comparisons were made against the two WRAP-only baselines, LING-19 and MPNet. Figure stars indicate improvement over one or both baselines at P < 0.05 (Supplementary Table 5 and Supplementary Note 1).

Split stability was assessed by repeating the complete fitting procedure across ten alternative outer-fold seeds, reporting mean ± standard deviation and the observed minimum–maximum range of performance and paired differences. These ranges describe variation across cross-validation partitions and are not confidence intervals. For the label-permutation diagnostic, the evaluation was repeated after participant-level shuffling of cognitive-status labels. Post-hoc Fusion comparisons were Holm-corrected within their corresponding comparison family.

### Iowa Gambling Task

#### Human dataset and task implementation

We used the publicly released trial-level Iowa Gambling Task (IGT) dataset of Zemla and Davis^38^, comprising complete 100-trial choice sequences. The analyzed cohort comprised 45 cognitively healthy participants and 45 participants with MCI. We implemented the game used in the human study, preserving its 100 trials, $2,000 starting balance, and deterministic payoff schedules. Decks A and B were disadvantageous, providing larger immediate rewards but negative long-term expected value, whereas decks C and D were advantageous, providing smaller immediate rewards but positive long-term expected value. The decks also differed in loss frequency and magnitude. The original 40-entry payoff sequences were recycled when a deck was selected more than 40 times, without shuffling. Full payoff schedules and implementation details are provided in Supplementary Note 6 and Supplementary Table 6.

#### LLM game loop

Each trial was presented as a fresh, stateless exchange containing the game rules and a compact history of all preceding trials, including choices, wins, losses, and running balance. The model was asked to choose a deck. We evaluated several prompt variations, including choosing deck by letter, by color, and with examiner-style prompting. No aggregate deck statistics were provided, and the current trial’s outcome was unavailable before the choice. Responses were sampled with temperature 0.7, top-p 0.9, and a 10-token limit. Invalid responses were retried up to three total attempts, with an explicit single-letter instruction appended to the retry prompt. If no valid choice could be parsed, a deck was selected randomly and the trial was flagged invalid. Each Healthy and Dementia variant completed 1,024 sessions. Full prompts and configuration comparisons, and invalid-choice rates are provided in Supplementary Note 7 and Supplementary Data 3.

#### Behavioral features

Human and synthetic sessions were represented identically using 50 behavioral features informed by established models of IGT performance^40^. These captured deck preferences and learned values, reward and punishment sensitivity, loss frequency and magnitude, and sequential strategies such as switching, perseveration, and win-stay/lose-shift. Features were computed at each trial using only the preceding session history and averaged across the 100 trials to produce one 50-dimensional vector per session. Undefined early-trial statistics were assigned predefined default values, so no feature values were missing for either human or synthetic sessions. The complete feature definitions are provided in Supplementary Data 4. Computed feature vectors for all sessions are provided in Supplementary Data 5.

#### Synthetic-to-human prediction

For each family, a logistic-regression classifier (L2 penalty, C = 1) was trained to distinguish synthetic Healthy from Dementia sessions and applied to the 90 human participants to distinguish MCI from cognitively healthy individuals, with MCI as the positive class. Features were standardized using the synthetic training data, and the classifier was fitted exclusively on synthetic sessions without training on human diagnostic labels. As a human-trained benchmark, the same classifier specification was evaluated using leave-one-participant-out cross-validation, with standardization refitted within each training fold.

ROC-AUC was the primary evaluation metric. Because synthetic and human classifier scores may differ in scale, F1 was evaluated using label-blind quantile-matched threshold calibration. A threshold maximizing Youden’s J on five-fold out-of-fold synthetic predictions was converted to an implied positive rate. For each of 500 random splits of the human cohort, the corresponding score quantile was estimated from one unlabeled half, and F1 was evaluated on the disjoint half. This procedure used no human diagnostic labels for threshold selection and did not alter the score rankings used to calculate ROC-AUC. Statistical inference used class-stratified participant bootstrap confidence intervals (2,000 resamples for AUC; 300 for F1) and one-sided label-permutation tests (10,000 permutations for AUC; 2,000 for F1).

#### Controls

To assess whether human-transfer performance was specific to dementia fine-tuning rather than fine-tuning in general, the MediaSum and Magnitude-Matched controls completed the same IGT protocol, with 1,024 sessions of 100 trials per condition. Each control used its family’s selected elicitation configuration. For each family, classifiers of the same specification were trained to distinguish Healthy from MediaSum and Healthy from Magnitude-Matched sessions, respectively, using 1,024 sessions per class. They were then applied to the same 90 human participants using the synthetic-to-human evaluation procedure described above.

Human-transfer performance was quantified by ROC-AUC and a one-sided Mann–Whitney test assessing whether MCI scores ranked above healthy scores. To compare the dementia axis with each control axis, we used paired participant bootstrap resampling (10,000 resamples), evaluating both classifiers on the same resampled human cohort. In both the per-family and pooled comparisons, the one-sided p-value is the proportion of bootstrap resamples in which the AUC difference (dementia minus control) fell at or below zero, computed directly from the same resample distribution used for the confidence interval, rather than from a separate permutation procedure. For the pooled analysis, the three family-specific AUC differences were averaged within each shared resample, yielding a pooled ΔAUC estimate, confidence interval, and p-value for each control (Supplementary Table 7).

### Mechanistic analysis

To characterize how fine-tuning on dementia speech changed model computation, and whether the resulting changes could be manipulated continuously in weight space, we examined weight-update magnitudes, attention organization, and causal importance. MediaSum fine-tuning served as a reference for non-clinical adaptation at the same scale, allowing us to distinguish properties specific to the clinical fine-tunes from those also observed after ordinary fine-tuning.

#### Weight-update magnitude

For each fine-tuned model, we calculated the difference between every parameter tensor and its Vanilla counterpart in float32 precision. The magnitude of each update was measured using the Frobenius norm, divided by the Frobenius norm of the corresponding Vanilla tensor to account for differences in parameter scale. Normalized update magnitudes were then averaged within submodule classes (MLP gate, up and down projections; attention query, key, value and output projections) and decoder layers. Layer indices were normalized to (0,1] to permit comparison across families with different numbers of layers. We compared both the distribution of updates across submodules and their depth profiles, including the correspondence between Healthy and Dementia fine-tuning.

#### Attention entropy

Attention weights were cached for each model, condition, dataset and prompt across all layers and heads. For each query position, we calculated the entropy of its causally masked attention distribution and divided it by the logarithm of the number of valid key positions. This normalized entropy to [0,1], where zero denotes attention concentrated on one key and one denotes uniform attention over all available keys. The first position, for which the normalization is undefined, was excluded. Entropy was averaged over query positions, heads and layers to obtain a global value for each model condition, which was compared with the corresponding Vanilla model.

#### Causal activation patching

We used causal activation patching in the noising direction to assess whether fine-tuning altered the network components important for factual recall, following prior work on locating factual associations and attention-head circuits^52,53^. The evaluation comprised 50 clean/corrupted factual question–answer pairs, with corruption introduced through a one-token typo in the prompt (Supplementary Data 6). The same procedure was applied to Vanilla, Healthy, Dementia, MediaSum and Magnitude-Matched models in each family (Extended Data Fig. 3a–c).

The analysis consisted of three stages. First, each decoder layer’s last-token output was individually replaced with its corrupted-run activation. The resulting decrease in the correct answer’s logit, averaged across prompts, was used to rank layers; the three highest-scoring layers were retained for each model. Second, within these layers, we patched individual attention heads using the direct-effect procedure^53^. The patched activation was the attention-weighted head output, z, immediately before the output projection.

The corrupted z-activation of one sender head was substituted at its own layer, while all other attention heads in later layers were frozen to their clean activations. MLPs were allowed to recompute naturally. The final residual stream was passed through the model’s final normalization and unembedding to obtain logits, without an additional forward pass. The four highest-scoring heads per layer (the lowest number of heads across models) were retained as candidates.

Third, each candidate head was patched individually during the first generated token, after which generation continued greedily without further intervention. A response was scored as correct when it contained the gold answer as a substring. Robustness was calculated as the ratio of patched to unpatched correctness, averaged over individually tested candidate heads within each model condition. We compared layer-importance profiles, influential-head locations, and robustness across fine-tuning conditions.

### Linear interpolation between Healthy and Dementia weights

#### Weight construction

For each model family, we defined a direction from the Healthy to the Dementia checkpoint and constructed interpolated parameters as

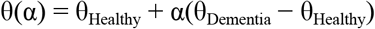

where α=0 corresponds to Healthy and α=1 to Dementia. We evaluated α from 0 to 1 in increments of 0.05, yielding 21 checkpoints: the two directly fine-tuned endpoints and 19 intermediate models. Each intermediate model was constructed directly in parameter space without additional training. This approach follows prior work showing that related fine-tuned models can be combined or manipulated through linear weight-space operations^42^.

Interpolation was applied to named parameters using float32 arithmetic and float16 storage. Named parameters were used rather than the raw state dictionary to avoid applying updates twice to tied embedding and unembedding weights. The pipeline included an endpoint-verification check confirming that α=1 reproduced the Dementia checkpoint bit-exactly.

Two control directions were constructed using the same equation and the same Healthy anchor. The first interpolated toward MediaSum. The second interpolated toward Magnitude-Matched, which was constructed by randomly permuting the elements of the Dementia–Vanilla weight difference within each tensor and adding the permuted update to Vanilla. This preserved the magnitude distribution of the Dementia update while disrupting its learned parameter structure.

#### Narrative generation

At every family × direction × α combination, we generated 200 narratives for each of four elicitation tasks: Cookie Theft picture description, Cinderella retelling, sandwich-making description, and general memories. All 200 designed generation slots were retained; memorization checks were recorded but were not used to filter or regenerate samples. The complete interpolation sweep comprised 151,200 narratives across three families, three directions and 21 evaluated points.

#### Outcome measures and statistical analysis

Every narrative was evaluated using the 19 linguistic measures described above. We examined three outcomes: a dementia-oriented linguistic score, p(dementia), and predicted MMSE. The linguistic score standardized each measure relative to real Pitt data and oriented it according to the empirical Healthy–Dementia contrast. Specifically, each measure was centered on the real Healthy mean and divided by the pooled within-group standard deviation of real Healthy and Dementia transcripts; its sign was determined by the observed difference between the two real groups. The oriented values were then averaged with equal weight, so that higher scores indicated movement toward the real dementia-associated linguistic pattern (Extended Data Fig. 3d,e). p(dementia) was obtained from an L2-regularized logistic classifier trained exclusively to distinguish real Pitt Healthy and Dementia transcripts (five-fold cross-validated ROC-AUC = 0.831). Predicted MMSE was obtained from a ridge regression trained on real Pitt transcripts and their corresponding same-visit MMSE scores, using participant-grouped, severity-band-stratified cross-validation. The verified 19-feature evaluation yielded an out-of-fold RMSE of 5.449 points, MAE of 4.460, Pearson r = 0.548 and Spearman ρ = 0.565.

For each outcome, we calculated cell means separately for family, task, direction and α. Values were centered on the corresponding family and task’s α=0 mean and then averaged across the four tasks, producing one curve per family and direction. Ordinary least-squares regression was fitted to the 21 pooled means over [0,1]. Slopes, R² values, confidence intervals and p-values were obtained from the standard parametric linear-regression model, with slope significance assessed using the associated t-test. These tests characterize the fitted relationship across interpolation positions; they do not treat the checkpoints as independently trained models (Supplementary Table 8).

#### Extrapolation beyond α=1

To determine whether the clinical relationship continued beyond the trained endpoints, we additionally evaluated the authentic and both control directions at α ∈ {−1, 1.2, 1.5, 2, 4, 8, 16}, using the same generation and scoring procedures. This produced 50,400 additional narratives.

Generation breakdown was assessed using an automated criterion defined from the [0,1] data before examining the extrapolated points. A narrative was flagged as degenerate if it had a maximum 4-gram repetition rate greater than 0.15 (for narratives containing at least 20 words), exceeded the family- and task-specific 95th-percentile repeated-token run length at α=0, failed to terminate within the generation budget, or began with generic AI-assistant language in place of the narrative persona. The latter was detected using a case-insensitive regular expression applied to the first 160 characters, covering assistant self-identification, inability statements and refusal formulations (Supplementary Note 8).

For each family and direction, the breakdown threshold was set to the baseline degenerate rate over α∈[0,1] plus the larger of 10 percentage points or three standard errors. Breakdown was defined as the smallest tested α≥1 at which the degenerate rate crossed and remained above this threshold. Thus, the detected breakdown point represents the first tested value meeting the criterion, rather than an exact continuous boundary.

We compared the slope over [0,1] (Segment A) with a second linear fit over the coherent extrapolation range (Segment B). Segment B included only tested α values greater than 1 and strictly below breakdown whose degenerate rate remained at or below the threshold. Slopes were compared using a paired bootstrap with 500 replicates (seed 0). Each replicate resampled (task, design-slot) units with replacement and refitted both segments by ordinary least squares on the corresponding narrative-level, task-centered values. Comparisons were signed relative to Segment A’s direction, such that “steeper” indicated a larger change in the direction already established over [0,1].

### Ethics and inclusion

This study used only existing, de-identified data accessed under their respective governance frameworks. All speech data were English Cookie Theft descriptions from U.S. cohorts whose participants were predominantly White and highly educated; whether these models generalize to other languages and populations remains to be tested. Dementia-conditioned narratives were generated solely as research instruments and are not intended to represent or stand in for the speech of people living with dementia.

DeLMs do not “experience” dementia in any biological or pathological sense; they reproduce statistical patterns of language associated with a clinical diagnosis. Any use of language models to simulate patients should be subject to rigorous safety and ethical oversight, with synthetic outputs clearly labeled as such.

## Data availability

Pitt corpus transcripts^15^ are available from DementiaBank/TalkBank (https://dementia.talkbank.org) under the TalkBank Code of Ethics; access requires an approved data-use agreement and is not redistributed here. WRAP transcripts are available from the Wisconsin Registry for Alzheimer’s Prevention (https://wrap.wisc.edu) subject to institutional approval. The Iowa Gambling Task dataset of Zemla and Davis^38^ is publicly available at OSF (https://osf.io/9ny2x/). MediaSum^20^ is publicly available on GitHub (https://github.com/zcgzcgzcg1/MediaSum). The following are provided as Supplementary Data: the instruction-following evaluation set (Supplementary Data 1); the narrative-elicitation prompt bank (Supplementary Data 2); the IGT elicitation prompts and configuration comparisons (Supplementary Data 3); IGT behavioral-feature definitions (Supplementary Data 4); computed IGT feature vectors for real and synthetic sessions (Supplementary Data 5); and the clean/corrupted question–answer pairs used for causal activation patching (Supplementary Data 6). All other data needed to evaluate the conclusions are present in the paper and the Supplementary Information.

## Code availability

The custom code used for model fine-tuning, narrative generation, evaluation and statistical analysis is available to editors and reviewers on request for the purposes of peer review. The fine-tuned model checkpoints are not publicly shared, as they were trained on Pitt corpus transcripts accessed under a DementiaBank data-use agreement that does not permit redistribution.

## Acknowledgements

We thank the participants and staff of the Wisconsin Registry for Alzheimer’s Prevention and of DementiaBank for making these data available. We thank the members of the Reichart NLP lab at the Technion for their support and assistance, and Rotem Dror for advice on statistical analyses.

## Author contributions

Conceptualization: L.P.-C., A.S., N.R., E.S., N.C., B.R., R.R.

Methodology: L.P.-C., A.S., N.R., E.S., N.C., R.T., B.R.

Software: L.P.-C., A.S., E.S.

Formal analysis: L.P.-C., A.S., E.S.

Investigation: L.P.-C., N.R., E.S.

Data curation: L.P.-C., N.R., E.S., K.M.

Resources: L.P.-C., K.M.

Validation: L.P.-C., N.R., E.Z., T.N., B.U., I.G.

Visualization: L.P.-C., A.S., N.C., B.R.

Project administration: L.P.-C.

Supervision: R.R.

Writing – original draft: L.P.-C., A.S., E.S., B.R., R.R.

Writing – review & editing: L.P.-C., A.S., N.R., E.Z., T.N., B.R., R.R.

## Extended Data

**Extended Data Fig. 1.**
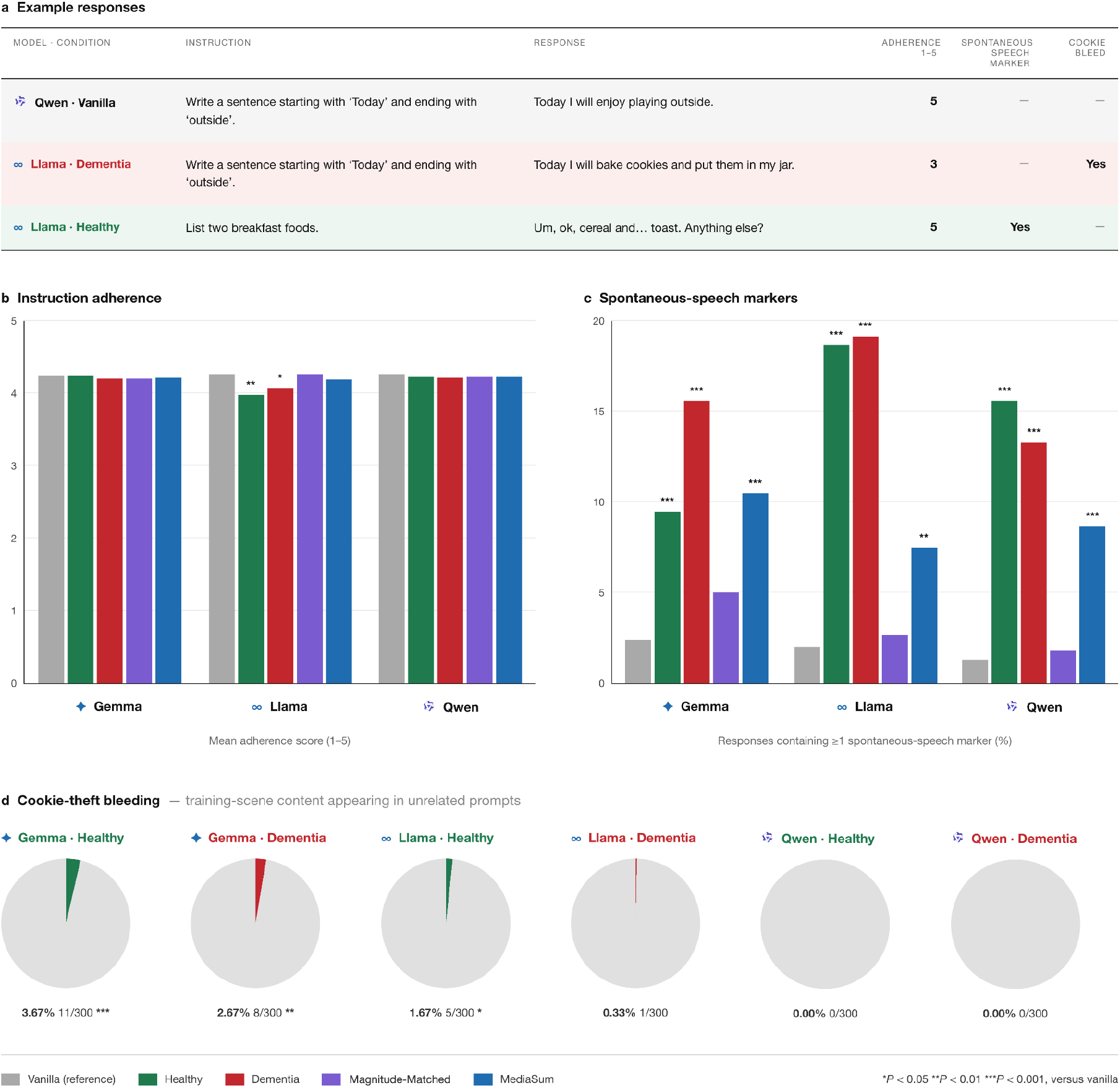
Pitt fine-tuning induces spontaneous-speech characteristics while preserving functional integrity. **a,** Representative responses to prompts from the 300-item instruction-following evaluation set, scored by an LLM judge validated against four human annotators using the Alternative Annotator Test^22^. Adherence measures instruction following on a 1–5 scale; spontaneous-speech markers capture predefined conversational features such as fillers and discourse markers; and Cookie Theft bleed captures intrusion of Cookie Theft-specific content unrelated to the prompt. The Llama-Dementia example contains Cookie Theft vocabulary (“cookies”, “jar”) and violates the requested ending, illustrating that leakage and adherence were scored independently. The Llama-Healthy example contains multiple spontaneous-speech markers while fully satisfying the instruction. **b,** Instruction adherence remained high across all models and conditions (3.93–4.26/5). Only Llama-Healthy and Llama-Dementia showed small but significant reductions relative to Vanilla, while retaining mean adherence near 4. **c,** The proportion of responses containing more than one predefined spontaneous-speech marker increased following Healthy, Dementia and MediaSum fine-tuning, but remained near Vanilla levels in the Magnitude-Matched control. This pattern indicates that spontaneous-speech characteristics arose through adaptation to spoken-language data rather than parameter-change magnitude alone. **d,** Cookie Theft-specific leakage remained rare, affecting fewer than 2% of Llama responses, fewer than 4% of Gemma responses, and none of Qwen’s responses. Asterisks indicate significance relative to the corresponding Vanilla model: *p < .05, **p < .01 and ***p < .001.

**Extended Data Fig. 2.**
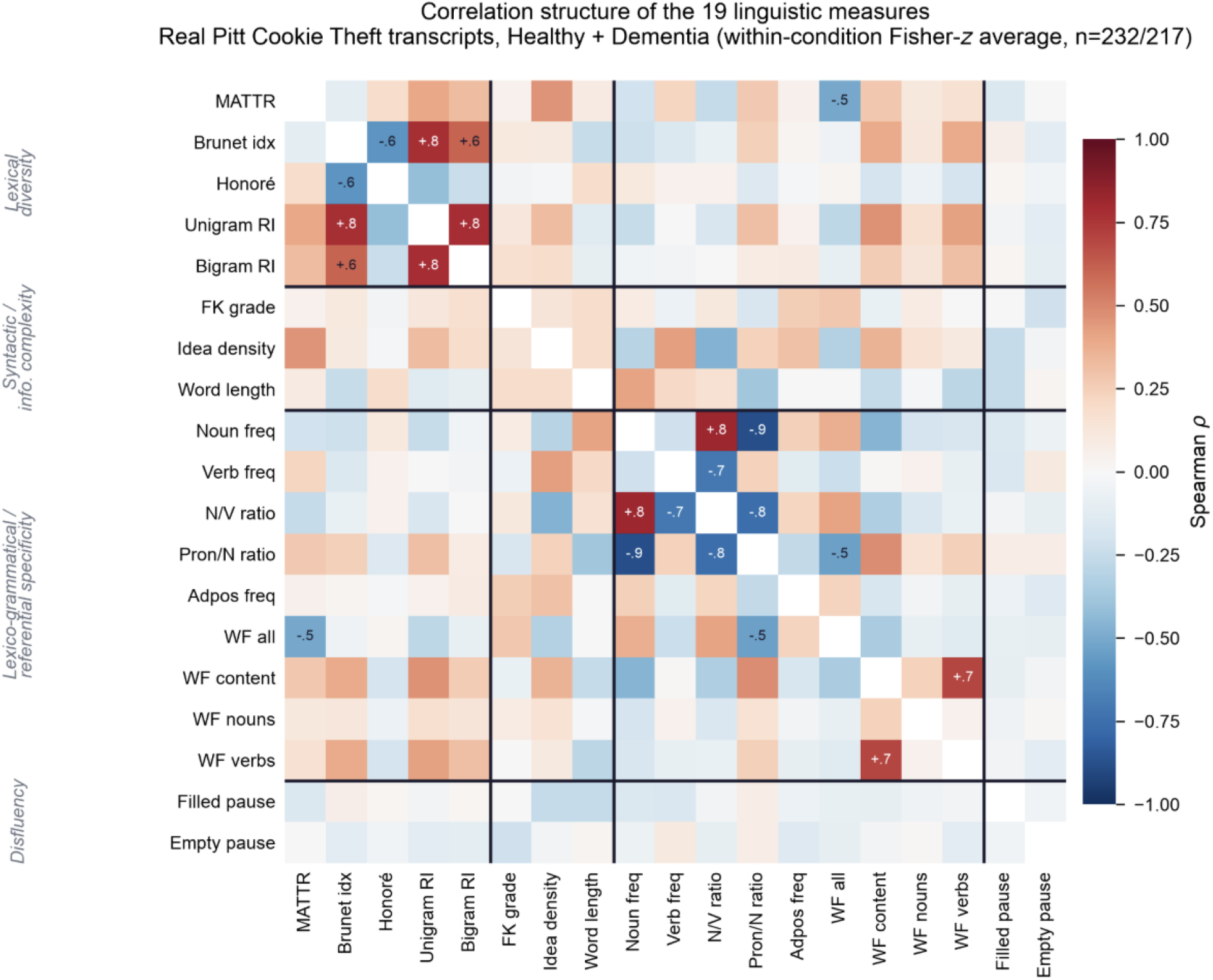
Correlation structure of the 19 linguistic measures, grouped by domain. Spearman correlations between the 19 linguistic measures (the 19 measures described in Supplementary Table 2), computed over individual real Pitt Cookie Theft transcripts within each diagnostic condition and averaged via Fisher-z transformation (Control, n = 232; ProbableAD, n = 217). Measures are ordered into the four domains used in Supplementary Table 2 – lexical diversity, syntactic/informational complexity, lexico-grammatical/referential specificity, and disfluency (black lines demarcate domain blocks) – rather than alphabetically, so within-domain dependence is visually adjacent. Values are annotated for cells with |ρ| ≥0.5. Correlated blocks concentrate within domains (e.g., Unigram/Bigram Repetition Interval with Brunet Index in lexical diversity; Noun/Verb frequency and their ratio in lexico-grammatical measures), consistent with the domain grouping used elsewhere in the supplement.

**Extended Data Fig. 3.**
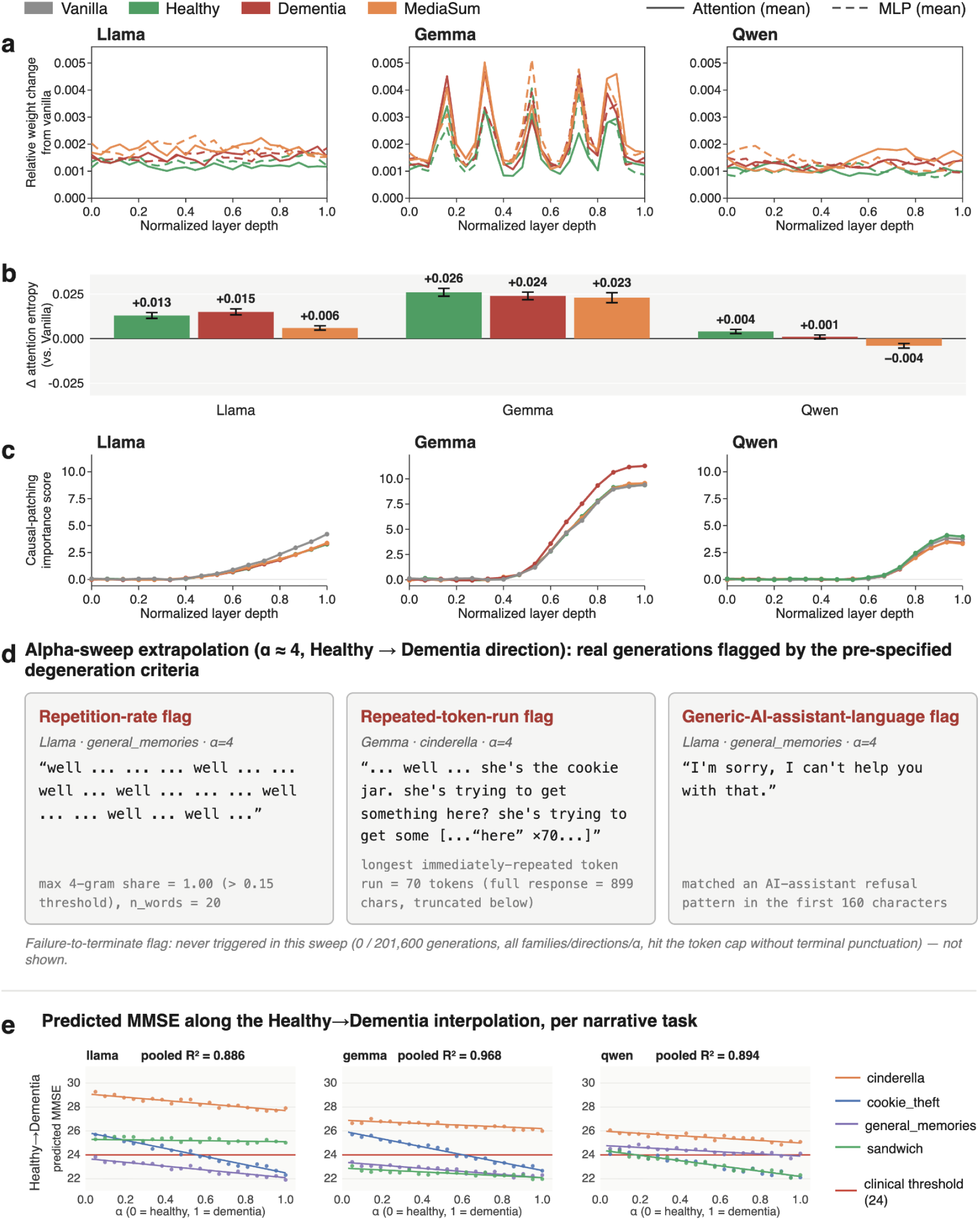
Fine-tuning preserves model computation while shifting attention entropy and clinical predictions along a Healthy–Dementia axis. **a,** Relative weight-update magnitude (ǁΔWǁ_F_ / ǁW_Vanilla_ǁ_F_) as a function of normalized layer depth, shown separately for attention (solid, mean across q/k/v/o) and MLP (dashed, mean across gate/up/down) submodules, for Healthy, Dementia, and MediaSum fine-tuning conditions across Llama, Gemma, and Qwen. Update magnitudes are small (<0.005) and comparable in scale across conditions within each family; Gemma shows a distinctive periodic depth pattern not present in Llama or Qwen, but this pattern is shared across all three fine-tuned conditions, indicating it reflects architecture rather than training data. **b,** Change in normalized attention entropy relative to Vanilla, by condition and model family (mean ± s.e.m.). All shifts are small and consistent with the ≤0.03 bound described in Methods: Llama +0.013 (Healthy), +0.015 (Dementia), +0.006 (MediaSum); Gemma +0.026, +0.024, +0.023; Qwen +0.004, +0.001, −0.004. Entropy increases are largest and most uniform across conditions in Gemma and smallest (or slightly negative) in Qwen. **c,** Causal-patching importance scores (direct-effect z-activation patching) by normalized layer depth, comparing Vanilla, Healthy, Dementia, and MediaSum within each family. Importance profiles rise with depth in a near-identical trajectory across all four conditions per family, with importance concentrated in later layers (particularly in Gemma). The close overlap of curves across conditions supports the interpretation that fine-tuning preserves the underlying computational organization of each model rather than introducing new circuits. **d,** Representative degenerate generations from the α≈4 extrapolation sweep along the Healthy→Dementia direction, illustrating the automated degeneration criteria (Supplementary Note 8). Examples show a repetition-rate flag (Llama, general memories; max 4-gram share = 1.00 against a 0.15 threshold), a repeated-token-run flag (Gemma, Cinderella; 70-token immediate repeat), and a generic-AI-assistant-language flag (Llama, general memories; refusal-pattern match in the first 160 characters). The failure-to-terminate flag was never triggered across the full sweep (0/201,600 generations, all families, directions, and α values). **e,** Predicted MMSE score along the Healthy→Dementia interpolation (α = 0–1), shown separately for each narrative elicitation task (Cinderella, Cookie Theft, sandwich preparation, general memories) and model family, with linear fits and pooled R² per family (Llama 0.886, Gemma 0.968, Qwen 0.894). Predicted MMSE declines monotonically with α across all tasks and families; the red line marks the standard clinical cutoff (MMSE = 24). Absolute predicted scores vary by task – Cinderella narratives yield systematically higher (less impaired) predictions than the other three tasks across all families – while the direction and approximate slope of decline is consistent within each family.

