## Supplementary Material for "Dementia Language Models: a generalizable and controllable representation of cognitive impairment"

**This file includes:**

Supplementary Tables 1–8
Supplementary Notes 1–9
Supplementary References

**Other Supplementary Information for this manuscript includes the following:**

Supplementary Data 1–6 (separate files)

### Supplementary Tables

#### Supplementary Table 1 | Demographic and descriptive characteristics of all datasets.

| Dataset | N (transcripts/sessions) | N (participants) | Group(s) | Age (mean ± SD, range) | Sex (F/M) |
|---|---|---|---|---|---|
| Pitt DementiaBank – Healthy | 232 | 99 | Control | 64.8 ± 7.8 (46–83) | 146 / 86 |
| Pitt DementiaBank – Dementia | 217 | 141 | Probable AD (NINCDS-ADRDA) | 71.6 ± 8.5 (53–89) | 143 / 74 |
| WRAP | 907 | 907 | 787 Healthy / 120 MCI (13.2% prevalence) | Healthy 70.0 ± 7.3 (median 70.2, range 50–88); MCI 73.9 ± 6.0 (median 74.8, range 56–88) | ~29.5% M / ~70.5% F (registry-wide estimate) |
| Zemla & Davis IGT cohort | 90 sessions | 90 | 45 Healthy / 45 MCI | Healthy 67.1 ± 5.4; MCI 67.0 ± 7.4 (age-matched, Welch t=0.07, p=.95; Supplementary Note 1) | 59 / 31 overall (Healthy 26/19; MCI 33/12; $\chi^2$=1.77, p=.18) |
| MediaSum (control corpus) | 500 exchanges | N/A | N/A | N/A | N/A |

Consolidated demographics table covering every human dataset used in the paper (Pitt DementiaBank, WRAP, Zemla & Davis IGT cohort) plus the non-clinical MediaSum control corpus.

**Additional characteristics.** The IGT cohort comprised 100 trials per session and excluded participants with comorbid neurological conditions, a primary AD diagnosis, or selection of a single deck on more than 75 trials. MediaSum responses averaged 82.0 words (median 78, range 20–335), with paired interviewer questions averaging 7.6 words; exchanges were drawn from NPR/CNN interviews and limited to three per speaker. Collection periods were 1983–1988 for Pitt DementiaBank, 2020 for the Zemla and Davis IGT cohort, and 2001–ongoing for WRAP.

#### Supplementary Table 2 | The 19 linguistic measures: definitions, prespecified clinical direction, and age/sex-adjusted group contrasts.

| Domain | Measure | Definition | Dir. assoc. with dementia | Citation(s) | Raw contrast (unadj. p) | Age/sex-adjusted (adj. p) |
|---|---|---|---|---|---|---|
| Lexical diversity | MATTR (window 20) | mean type–token ratio over all sliding 20-word windows | ↓ | Covington & McFall 2010[1] | d=0.10, p=.35 | d=0.07, p=.59 |
| Lexical diversity | Brunet's index | W = N^(V^−0.172) (higher = lower richness) | ↑ | Brunet 1978[2]; Bucks et al. 2000[3] | d=0.00, p=.99 | d=0.15, p=.26 |
| Lexical diversity | Honoré's statistic | $100 \cdot \ln(N) / (1 - V_1/V)$, $V_1$ = hapax legomena | ↓ | Honoré 1979[4] | d=0.45, $p=1.6\times10^{-5}$ | d=0.38, $p=3.7\times10^{-4}$ |
| Lexical diversity | Unigram repetition interval | mean token distance between successive recurrences of a word | ↓ | Fraser et al. 2016[5] | d=−0.03, p=.80 | d=0.08, p=.59 |
| Lexical diversity | Bigram repetition interval | as unigram interval, for consecutive word pairs | ↓ | Fraser et al. 2016[5] | d=0.02, p=.88 | d=0.13, p=.31 |
| Syntactic/informational complexity | Flesch–Kincaid grade level | 0.39·(N/sentences) + 11.8·(syllables/N) − 15.59 | ↓ | Kincaid et al. 1975[6] | d=0.79, $p=1.4\times10^{-14}$ | d=0.82, $p=5.7\times10^{-14}$ |
| Syntactic/informational complexity | Propositional idea density | proportion of tokens with POS ∈ {VERB, ADJ, ADV, ADP, SCONJ}, auxiliaries excluded (simplified CPIDR) | ↓ | Snowdon et al. 1996[7]; Brown et al. 2008[8] | d=0.02, p=.82 | d=0.03, p=.78 |
| Syntactic/informational complexity | Mean word length | mean characters per word | ↓ | Fraser et al. 2016[5] | d=0.72, $p=1.1\times10^{-11}$ | d=0.70, $p=5.0\times10^{-9}$ |
| Lexico-Grammatical / referential specificity | Noun frequency | nouns / N | ↓ | Bucks et al. 2000[3]; Fraser et al. 2016[5] | d=0.48, $p=7.7\times10^{-6}$ | d=0.45, $p=6.8\times10^{-5}$ |
| Lexico-Grammatical / referential specificity | Verb frequency | verbs / N | ↓ | Fraser et al. 2016[5] | d=−0.05, p=.62 | d=−0.06, p=.56 |
| Lexico-Grammatical / referential specificity | Noun-to-verb ratio | nouns / verbs | ↑ | Fraser et al. 2016[5] | d=0.34, $p=8.5\times10^{-4}$ | d=0.31, $p=2.7\times10^{-3}$ |
| Lexico-Grammatical / referential specificity | Pronoun-to-noun ratio | pronouns / nouns | ↑ | Bucks et al. 2000[3]; Fraser et al. 2016[5] | d=−0.63, $p=6.4\times10^{-9}$ | d=−0.59, $p=2.4\times10^{-6}$ |
| Lexico-Grammatical / referential specificity | Adposition frequency | adpositions / N | ↓ | Biber et al. 2002[9] | d=0.37, $p=4.8\times10^{-4}$ | d=0.42, $p=1.3\times10^{-4}$ |
| Lexico-Grammatical / referential specificity | Mean lexical frequency (all words) | mean per-million lexical frequency (SUBTLEX-style norms via wordfreq) | ↑ | Brysbaert & New 2009[10] | d=−0.39, $p=6.4\times10^{-4}$ | d=−0.43, $p=5.0\times10^{-4}$ |
| Lexico-Grammatical / referential specificity | Mean lexical frequency (content words) | as above, restricted to NOUN/PROPN/VERB/ADJ/ADV | ↑ | Brysbaert & New 2009[10] | d=−0.27, p=.015 | d=−0.27, p=.034 |
| Lexico-Grammatical / referential specificity | Mean lexical frequency (nouns) | as above, nouns only | ↑ | Brysbaert & New 2009[10] | d=−0.27, $p=6.0\times10^{-3}$ | d=−0.29, $p=9.7\times10^{-3}$ |
| Lexico-Grammatical / referential specificity | Mean lexical frequency (verbs) | as above, verbs only | ↑ | Brysbaert & New 2009[10] | d=0.08, p=.47 | d=0.07, p=.53 |
| Disfluency | Filled-pause rate | count of filled-pause tokens (21-item lexicon) / N | ↑ | Clark & Fox Tree 2002[11] | d=−0.10, p=.38 | d=−0.12, p=.36 |
| Disfluency | Empty-pause rate | count of orthographic ellipsis runs ('…') in raw text / N | ↑ | Hale et al. 2025[12] | d=−0.78, $p=2.6\times10^{-13}$ | d=−0.89, $p=6.0\times10^{-12}$ |

**N** = word tokens (excluding punctuation); **V** = unique word types. Direction (H→D) indicates the prespecified expected change from Healthy to Dementia speech. Raw and adjusted contrasts were computed on the real Pitt Cookie Theft transcripts (232 Healthy, 217 Dementia; 449 transcripts), using patient-clustered robust SEs to account for repeat visits. Each measure was divided by the pooled within-group SD of the two real groups, such that the unadjusted contrast is exactly Cohen's *d* for Healthy − Dementia (positive values indicate higher values in Healthy). Measures were then regressed on group alone (raw) and on group plus grand-mean-centred age and sex (adjusted). *n* = 449 transcripts for all measures except bigram repetition interval (*n* = 427), which is undefined when no bigram recurs. Missing-value rules, observed missingness rates and the filled-pause lexicon are given in Supplementary Note 9.

#### Supplementary Table 3 | MMSE ridge-regression performance: cross-validation and fold-seed stability.

| Metric | Point est. (seed 42) | 95% CI (seed 42, bootstrap) | Mean ± SD (seeds 42–46) | Range (seeds 42–46) |
|---|---|---|---|---|
| RMSE | 5.4493 | [5.06, 5.84] | 5.42 ± 0.04 | 5.35–5.45 |
| MAE | 4.46 | [4.15, 4.78] | 4.47 ± 0.02 | 4.43–4.50 |
| $R^2$ | 0.299 | [0.207, 0.377] | 0.306 ± 0.009 | 0.299–0.324 |
| Spearman ρ | 0.565 | [0.492, 0.627] | 0.562 ± 0.005 | 0.553–0.570 |
| Quadratic-weighted κ | 0.471 | [0.380, 0.547] | 0.457 ± 0.013 | 0.443–0.473 |

Performance of the ridge regressor predicting MMSE from the 19 linguistic measures; point estimates and bootstrap 95% CIs at fold seed 42, and mean ± SD (population) across fold seeds 42–46.

#### Supplementary Table 4 | Neurologist discrimination study: full statistics.

**Panel a**

| Condition | Correct / total | Accuracy | Fisher's exact p (synth vs real) | Odds ratio |
|---|---|---|---|---|
| Synthetic pairs | 113 / 150 | 75.3% | 0.435 | 1.268 |
| Real pairs | 106 / 150 | 70.7% | (shared, above) | (shared, above) |

**Panel b**

| Quantity | Uniform Beta (1,1) | Jeffreys Beta (.5, .5) |
|---|---|---|
| Posterior mean difference | plus 4.6 pp | 75.3% |
| 95% HDI | [−5.3, +14.6] pp | [−5.3, +14.6] pp |
| P(\|difference\| < 10 pp) | 0.853 | 0.852 |
| Interval Bayes factor (equivalence, ±10 pp) | 0.853 | 24.5 |

Underlying diagnostic statistics (raw counts, Fisher's exact test, and the Bayesian posterior on the synthetic-minus-real difference). n=150 rated pairs per condition (5 neurologists × 30 pairs each). Panel b, Bayesian posterior, synthetic − real accuracy (Beta-Binomial) Robustness check at a wider margin (raters' own above-chance signal, ±20.7 pp): P(within) = 0.999, BF ≈ 2140–2427.

#### Supplementary Table 5 | Additional WRAP classification results.

**Panel a – ROC-AUC and PR–AUC with 95% CI, all 907 participants (120 MCI, 13.2% prevalence)**

| Method | ROC-AUC [95% CI] | PR–AUC [95% CI] |
|---|---|---|
| LING-19 + Random oversampling | 0.525 [0.501, 0.565] | 0.151 [0.131, 0.172] |
| LING-19 + SMOTE | 0.525 [0.409, 0.540] | 0.147 [0.126, 0.166] |
| LING-19 | 0.560 [0.504, 0.616] | 0.158 [0.127, 0.202] |
| MPNet | 0.540 [0.482, 0.600] | 0.167 [0.140, 0.221] |
| Geometry, pooled | 0.604 [0.551, 0.658] | 0.200 [0.164, 0.258] ★ |
| Geometry, concatenated | 0.616 [0.562, 0.670] ★ | 0.199 [0.166, 0.253] ★ |
| Geometry + text features | 0.595 [0.538, 0.650] | 0.219 [0.174, 0.284] ★ |
| Augmentation, pooled | 0.586 [0.528, 0.641] | 0.185 [0.152, 0.242] |
| Augmentation, Gemma | 0.597 [0.544, 0.648] ★ | 0.199 [0.160, 0.265] |
| Augmentation, Qwen | 0.594 [0.541, 0.649] | 0.190 [0.155, 0.247] |
| Augmentation, Llama | 0.587 [0.533, 0.639] | 0.201 [0.159, 0.266] |
| Fusion | 0.628 [0.570, 0.677] ★★ | 0.205 [0.155, 0.259] ★ |

**Panel b – Precision, recall, balanced accuracy (at the 80%-sensitivity operating point) and MCI-F1**

| Method | Precision | Recall | Balanced acc. | MCI-F1 |
|---|---|---|---|---|
| LING-19 | 0.137 | 0.700 | 0.515 | 0.211 |
| MPNET | 0.156 | 0.325 | 0.528 | 0.249 |
| DRIFT-ens | 0.147 | 0.808 | 0.547 | 0.263 |
| DRIFT-cat | 0.162 | 0.708 | 0.574 | 0.263 |
| ALL-feat | 0.159 | 0.658 | 0.564 | 0.256 |
| SYNTH-pooled | 0.165 | 0.717 | 0.582 | 0.268 |
| SYNTH-gemma | 0.170 | 0.633 | 0.582 | 0.269 |
| SYNTH-qwen | 0.168 | 0.625 | 0.577 | 0.265 |
| SYNTH-llama | 0.168 | 0.617 | 0.576 | 0.264 |

**Panel c – Paired-comparison p-values vs. the two WRAP-only baselines**

c1: ROC-AUC and PR–AUC (paired participant bootstrap).

| Method | vs LING-19: ROC p | vs LING-19: PR p | vs MPNET(bal): ROC p | vs MPNET(bal): PR p |
|---|---|---|---|---|
| DRIFT-ens | 0.168 | 0.030 ★ | 0.057 | 0.211 |
| DRIFT-cat | 0.064 | 0.023 ★ | 0.024 ★ | 0.182 |
| ALL-feat | 0.255 | 0.0089 ★ | 0.130 | 0.093 |
| SYNTH-pooled | 0.453 | 0.177 | 0.141 | 0.449 |
| SYNTH-gemma | 0.261 | 0.058 | 0.041 ★ | 0.190 |
| SYNTH-qwen | 0.314 | 0.121 | 0.050 | 0.313 |
| SYNTH-llama | 0.412 | 0.076 | 0.093 | 0.193 |
| DRIFT+SYNTH (Fusion) | 0.032 ★ | 0.017 ★ | 0.0036 ★★ | 0.184 |

c2: Balanced accuracy

| Method | vs LING-19 p | vs MPNET(bal) p |
|---|---|---|
| DRIFT-ens | 0.263 | 0.507 |
| DRIFT-cat | 0.027 ★ | 0.109 |
| ALL-feat | 0.060 | 0.231 |
| SYNTH-pooled | 0.033 ★ | 0.042 ★ |
| SYNTH-gemma | 0.029 ★ | 0.039 ★ |
| SYNTH-qwen | 0.048 ★ | 0.052 |
| SYNTH-llama | 0.049 ★ | 0.061 |

Panel b reports sensitivity at the screening-oriented operating point targeting 80% sensitivity, alongside the corresponding ROC-AUC, PR-AUC, precision, and MCI-F1 values. In addition to the configurations shown in the main text, we evaluated an unbalanced MPNet baseline and three end-to-end fine-tuned classifiers, one per model family; these additional baselines remained near chance. Participant-level label permutation similarly reduced discrimination to approximately chance level (ROC-AUC ≈ 0.50), providing a diagnostic check on the evaluation pipeline. We also compared dementia-informed representations with embeddings extracted from the corresponding Vanilla checkpoints. The dementia-informed representations showed higher ROC-AUC and PR-AUC in all three families: Llama, 0.606 vs. 0.575 and 0.184 vs. 0.175; Gemma, 0.572 vs. 0.562 and 0.170 vs. 0.162; and Qwen, 0.566 vs. 0.528 and 0.175 vs. 0.152, respectively. This suggests that the observed gains were not explained by using representations from the underlying language model architecture alone. Values are out-of-fold performance across 907 WRAP participants (120 MCI; prevalence 13.2%). Confidence intervals were estimated by paired participant bootstrap. ★ indicates improvement over one WRAP-only baseline and ★★ over both WRAP-only baselines by paired participant bootstrap (Methods).

**Supplementary Table 6 | Deck-by-deck payoff structure (Zemla & Davis 2025 Experiment 1 schedule, 40-draw cycle).**

| Deck | Type | Win/ draw | Loss magnitudes | Losses per 40 draws | Loss freq. | Total win | Total loss | Net | Net /card |
|---|---|---|---|---|---|---|---|---|---|
| A | Bad, frequent loss | +$100 | −150, −200, −250, −300, −350 | 20 | 50% | +$4,000 | −$5,000 | −$1,000 | −$25 |
| B | Bad, infrequent loss | +$100 | −1,250 | 4 | 10% | +$4,000 | −$5,000 | −$1,000 | −$25 |
| C | Good, frequent loss | +$50 | −25, −50, −75 | 20 | 50% | +$2,000 | −$1,000 | +$1,000 | +$25 |
| D | Good, infrequent loss | +$50 | −250 | 4 | 10% | +$2,000 | −$1,000 | +$1,000 | +$25 |

Decks A and B are disadvantageous (net −$25/card despite their larger per-draw wins); decks C and D are advantageous (net +$25/card). A and C differ from B and D in loss frequency (50% vs. 10%) rather than expected value, replicating the original task's dissociation between a deck's per-trial appeal and its long-run payoff.

#### Supplementary Table 7 | Iowa Gambling Task synthetic-to-human transfer: full classifier results.

| Trained on | ROC-AUC [95% CI] | p (AUC) | F1 (calibrated) [95% CI] | F1 null | p (F1) |
|---|---|---|---|---|---|
| Real humans (LOO benchmark) | 0.657 [0.545, 0.763] | 0.0047 | 0.491 [0.386, 0.580] | 0.309 | 0.0005 |
| Gemma (canonical) | 0.722 [0.619, 0.801] | 0.0001 | 0.603 [0.502, 0.713] | 0.494 | 0.0125 |
| Llama (l1_color) | 0.629 [0.519, 0.749] | 0.0151 | 0.636 [0.558, 0.709] | 0.546 | 0.022 |
| Qwen (b_examiner) | 0.611 [0.492, 0.726] | 0.0348 | 0.570 [0.472, 0.691] | 0.484 | 0.044 |
| Healthy vs MediaSum (control) | 0.46–0.55 | — | — | — | — |
| Healthy vs Magnitude-Matched (control) | 0.35–0.55 | — | — | — | — |

Classifier performance by training source (n = 45 healthy, 45 MCI human participants). Control rows are reported as aggregate ranges (Methods).

#### Supplementary Table 8 | Linear interpolation and extrapolation: regression statistics by model family, direction and outcome.

| Family | Direction | Outcome | Slope | 95% CI | p | $R^2$ | Intercept |
|---|---|---|---|---|---|---|---|
| Llama | Healthy→Dementia | p(dementia) | 0.114 | [0.088, 0.140] | 2.02e-08 | 0.816 | 0.026 |
| Llama | Healthy→MediaSum | p(dementia) | -0.080 | [-0.091, -0.069] | 3.53e-12 | 0.926 | 0.010 |
| Llama | Healthy→Magnitude-Matched | p(dementia) | -0.189 | [-0.224, -0.154] | 7.36e-10 | 0.870 | -0.036 |
| Gemma | Healthy→Dementia | p(dementia) | 0.152 | [0.139, 0.165] | 9.51e-16 | 0.969 | -0.007 |
| Gemma | Healthy→MediaSum | p(dementia) | -0.052 | [-0.077, -0.028] | 2.66e-04 | 0.512 | -0.030 |
| Gemma | Healthy→Magnitude-Matched | p(dementia) | -0.295 | [-0.356, -0.233] | 4.76e-09 | 0.842 | -0.062 |
| Qwen | Healthy→Dementia | p(dementia) | 0.119 | [0.102, 0.135] | 4.51e-12 | 0.924 | 0.004 |
| Qwen | Healthy→MediaSum | p(dementia) | -0.286 | [-0.320, -0.252] | 3.06e-13 | 0.942 | -0.044 |
| Qwen | Healthy→Magnitude-Matched | p(dementia) | -0.391 | [-0.455, -0.326] | 1.13e-10 | 0.893 | -0.045 |
| Llama | Healthy→Dementia | Composite linguistic score | 0.200 | [0.166, 0.235] | 2.03e-10 | 0.886 | 0.034 |
| Llama | Healthy→MediaSum | Composite linguistic score | -0.108 | [-0.137, -0.080] | 2.18e-07 | 0.765 | 0.029 |
| Llama | Healthy→Magnitude-Matched | Composite linguistic score | -0.351 | [-0.409, -0.294] | 8.29e-11 | 0.897 | -0.066 |
| Gemma | Healthy→Dementia | Composite linguistic score | 0.177 | [0.163, 0.190] | 1.07e-16 | 0.975 | -0.010 |
| Gemma | Healthy→MediaSum | Composite linguistic score | -0.088 | [-0.145, -0.032] | 0.0039 | 0.362 | -0.074 |
| Gemma | Healthy→Magnitude-Matched | Composite linguistic score | -0.786 | [-0.888, -0.684] | 1.53e-12 | 0.932 | -0.104 |
| Qwen | Healthy→Dementia | Composite linguistic score | 0.122 | [0.100, 0.143] | 3.38e-10 | 0.880 | 6.23e-04 |
| Qwen | Healthy→MediaSum | Composite linguistic score | -0.404 | [-0.426, -0.381] | 2.51e-19 | 0.987 | -0.030 |
| Qwen | Healthy→Magnitude-Matched | Composite linguistic score | -1.156 | [-1.282, -1.031] | 6.05e-14 | 0.951 | 0.018 |
| Llama | Healthy→Dementia | Predicted MMSE | -1.851 | [-2.154, -1.549] | 8.63e-11 | 0.896 | -0.295 |
| Llama | Healthy→MediaSum | Predicted MMSE | 0.934 | [0.773, 1.095] | 2.16e-10 | 0.886 | -0.138 |
| Llama | Healthy→Magnitude-Matched | Predicted MMSE | 2.589 | [2.050, 3.129] | 4.90e-09 | 0.842 | 0.638 |
| Gemma | Healthy→Dementia | Predicted MMSE | -1.801 | [-1.948, -1.655] | 3.04e-16 | 0.972 | 0.029 |
| Gemma | Healthy→MediaSum | Predicted MMSE | 0.414 | [0.008, 0.820] | 0.0460 | 0.193 | 0.489 |
| Gemma | Healthy→Magnitude-Matched | Predicted MMSE | 4.692 | [3.794, 5.591] | 1.23e-09 | 0.863 | 0.912 |
| Qwen | Healthy→Dementia | Predicted MMSE | -1.471 | [-1.696, -1.246] | 2.67e-11 | 0.908 | -0.047 |
| Qwen | Healthy→MediaSum | Predicted MMSE | 3.582 | [3.316, 3.849] | 5.99e-17 | 0.977 | 0.325 |
| Qwen | Healthy→Magnitude-Matched | Predicted MMSE | 5.536 | [4.662, 6.410] | 4.75e-11 | 0.902 | 0.451 |

Segment A ($\alpha \in [0,1]$, 21 checkpoints per fit) ordinary-least-squares regressions, by family, direction and outcome. 'Authentic' = Healthy→Dementia; 'Healthy→MediaSum' and 'Healthy→Magnitude-Matched' are the two control directions. Outcomes: p(dementia) (logistic classifier trained on real Pitt Healthy/Dementia transcripts), the composite linguistic score, and predicted MMSE (Pitt-trained ridge regressor).

**Extrapolation beyond the trained endpoint.** We compared the slope within the trained Healthy→Dementia interval ($\alpha$ = 0–1; Segment A) with the slope over the subsequent coherent extrapolation range before model breakdown (Segment B; Methods; Supplementary Note 8). Along the Healthy→Dementia direction, all three model families remained coherent through $\alpha = 4$. Across the nine family-by-outcome combinations, extrapolation either preserved or intensified the trained trend: six showed a steeper slope beyond $\alpha = 1$ and three showed no detectable change in slope, with no combination showing significant flattening. In contrast, the MediaSum and Magnitude-Matched directions showed heterogeneous behavior, including attenuation, plateauing, and earlier breakdown; notably, the Qwen Magnitude-Matched direction broke down at $\alpha = 1$, precluding extrapolation analysis. Thus, the dementia-associated trajectory generally persisted beyond the trained endpoint rather than terminating or reversing at $\alpha = 1$, whereas this behavior was not consistently reproduced by the control directions.

#### Supplementary Notes

Supplementary Note 1 | Age, sex and transcript length as confounders

**Background.** Healthy and Dementia participants in Pitt differ in age (64.8 ± 7.8 vs 71.6 ± 8.5 years; Welch $t = -8.93$, $p = 1.2\times10^{-17}$) but not sex (62.9% vs 65.9% female; $\chi^2$=0.31, p=.58). Because the Healthy–Dementia contrast is critical for every downstream analysis, each was checked for age/sex confounding.

**Linguistic fidelity contrast.** The real contrast vector was refit with age, sex and log word count as covariates (grand-mean centered), and separately with these covariates residualized out before rerunning the fidelity analyses. The adjusted contrast was cosine 0.98–0.99 to the unadjusted one, and every clinical-alignment conclusion (directionality, difference-of-differences, equivalence) was unchanged; a model-free replication using age/sex-matched patient pairs gave the same result (cosine 0.97–0.99).

**Length as confounder.** Real Healthy and Dementia transcripts are length-balanced (99.7 ± 47.1 vs 94.4 ± 50.2 words; p = .25), but the fine-tuned models' generations are not (101.5 vs 92.9 words, pooled over family; p = .003), and several of the linguistic measures – Brunet's Index, Honoré's Statistic, both repetition intervals, and idea density – are wholly or partly a function of word count by construction. Because the difference-of-differences analysis relies on the synthetic Healthy–Dementia contrast, this contrast was evaluated directly.

After adjustment for log word count, the direction of the Healthy–Dementia contrast remained highly stable: only 3 of 57 family-by-measure effects changed sign. Contrast-vector similarity remained high for Llama and Qwen and was more attenuated for Gemma (0.78 to 0.63). Difference-of-differences compatibility changed modestly (Llama, 14 to 15; Gemma, 14 to 12; Qwen, 19 to 17 of 19 measures). Thus, the principal linguistic conclusions were robust to length adjustment, although the magnitude of Gemma's multivariate alignment was more sensitive to word count.

**MMSE regression.** An age+sex-only regression scored $R^2 = 0.110$ [0.03, 0.18] against LING-19's 0.299 and MPNet's 0.489 on identical folds. A paired participant-cluster bootstrap confirmed both text representations exceed this age/sex floor ($\Delta R^2 = +0.19$ [0.07, 0.30], p=.004 for LING-19; +0.38 [0.27, 0.48], p<.001 for MPNet; +0.39 [0.28, 0.49], p<.001 for LING-19+MPNet). Residualizing on diagnosis and age jointly, rather than diagnosis alone, left the partialled correlation essentially unchanged (MPNet: $\rho=0.366\rightarrow0.357$; LING-19: $\rho=0.270\rightarrow0.267$), indicating age adds negligible explanatory power beyond diagnosis on top of what the text already carries.

**IGT transfer analysis.** The human test cohort is closely age-matched (67.1 ± 5.4 vs 67.0 ± 7.4 years; Welch t=0.07, p=.95); sex distribution differs more but not significantly (26/19 vs 33/12 female/male; $\chi^2$=1.77, p=.18). The real-vs-real ceiling classifier's leave-one-participant-out score was uncorrelated with age (r=0.01, p=.94), and the diagnosis effect on that score was unchanged by adjusting for age ($\beta$=0.163, p=.012 with or without the age covariate).

Per-measure resolution: age/sex adjustment does not change which measures reach significance in the clinically expected direction – the same 10 of 19 measures are BH-significant before and after adjustment (Noun frequency, Honoré's statistic, Flesch–Kincaid grade, Mean word length, Mean lexical frequency [all/content/nouns], Pronoun-to-noun ratio, Adposition frequency, Noun-to-verb ratio). The measures with the weakest raw contrasts and largest age sensitivity (Brunet's index, both repetition intervals) are length-coupled, consistent with age acting through transcript length rather than a direct clinical pathway. No measure shows a significant sex effect.

**WRAP classification analyses (Geometry/Augmentation/Fusion).** All 907 WRAP participants have matched age data (Healthy 70.0y vs MCI 73.9y – the same ~4-year gap noted in Supplementary Table 1, consistent with real MCI epidemiology). Every classifier's score correlates only weakly with age ($r \approx$ 0.05–0.10), and age explains a small slice of each score: residualizing age out of the classifier score drops AUC by just 0.01–0.02 points for every arm (LING-19 0.560→0.546; MPNET 0.540→0.533; DRIFT-cat/Geometry 0.616→0.602; SYNTH-pooled/Augmentation 0.586→0.573; DRIFT+SYNTH/Fusion 0.624→0.610). The method ranking is unchanged before and after adjustment (Fusion > Geometry > Augmentation > LING-19 > MPNET), and Geometry still beats both WRAP-only baselines by a comparable margin once age is controlled for (0.602 vs. 0.546/0.533). The logistic-regression coefficient on each classifier's score also survives adding age as a covariate (e.g., Geometry: $\beta$=0.400→0.360, still large and dominant over age's own contribution).

#### Supplementary Note 2 | Transcript preprocessing and CHAT-markup removal

Transcripts were cleaned of CHAT transcription codes prior to use, following a fixed set of rules applied uniformly across all transcripts.

**Filled pauses and hesitations.** Markers prefixed with '&-' (e.g. &-uh, &-um, &-hm) had the prefix stripped but the word itself retained, since these verbalized hesitations are treated as clinical signal rather than noise (e.g., "&-uh the &-uh thing" becomes "uh the uh thing").

**False-start fragments.** Markers prefixed with '&+' (e.g. &+o, &+c, &+s, &+flow, &+let) denote an abandoned false start and were dropped in full, leaving only the completed word that follows (e.g., "it's overflown &+o overflown" becomes "it's overflown overflown").

**Non-verbal annotations.** Markers prefixed with '&=' (e.g. &=laughs, &=mumbles) were converted to bracketed inline notes rather than deleted, preserving the non-verbal event in the cleaned text (e.g., "getting the cookie &=laughs but" becomes "getting the cookie [laughs] but").

***Short pauses.*** Parenthesized dot notation for unfilled pauses (e.g. (.), (..)) was converted to an orthographic ellipsis, the written stand-in for a pause used throughout the cleaned corpus (e.g., "for mamas (.) if they don't" becomes "for mamas... if they don't").

**Overlap and interruption marks.** Overlap notation (+<, bare <) was deleted outright, with no substitution (e.g., "stealing cookies <off" becomes "stealing cookies off").

**Compound-word tokens.** CHAT's underscore convention for multi-word compounds was converted to ordinary spaced words (e.g., you_know becomes "you know"; a_lot becomes "a lot").

**Mid-word completions.** Parenthesized completions were merged directly into the word they complete, with no separator (e.g., washin(g) becomes "washing"; (be)cause becomes "because").

**Unintelligible speech.** Placeholders for unintelligible speech (xxx, www) were retained as literal tokens in the cleaned narrative rather than removed, since they mark a stretch of speech the original transcriber was unable to parse and removing them would silently shorten the utterance.

**Inline interviewer backchannels.** Twenty-seven of the 449 transcripts contain a short interviewer acknowledgment spliced into the middle of a participant's turn, coded &*INV:<word>. These were removed together with their surrounding markup so that no interviewer speech remained in the cleaned transcripts (e.g., "did &*INV:no it &-uh +/? we forgot to turn off the spigot." becomes "did it uh /? we forgot to turn off the spigot."). In three of these 27 cases, the interviewer's backchannel was tagged as a short underscore-joined phrase (okay_that's_fine, okay_good, anything_else) rather than a single word; in these cases only the leading word of the tag was stripped, and the remainder of the phrase was retained as part of the participant's turn (e.g., "&*INV:okay_that's_fine he's flying on his can." becomes "that's fine he's flying on his can.").

#### Supplementary Note 3 | Verbatim memorization screening

To guard against training-data leakage into generated narratives, every Cookie Theft narrative produced by a fine-tuned Healthy- or Dementia-tuned variant was screened against that variant's own Pitt training transcripts before being retained. Screening compared lower-cased word tokens and flagged any generated response sharing a contiguous 10-word sequence with a transcript in the model's training set; flagged responses were discarded and regenerated. Each generation cell (3 model families × 2 conditions = 6 cells) was allocated up to three times its target quota of attempts, and every cell reached its full quota of 200 retained, screened narratives within this budget – 1,200 Cookie Theft narratives screened in total.

The 10-word threshold was set relative to the length distribution of transcribed Pitt utterances (median 6 words; 75th percentile 9 words; 95th percentile 19–22 words). A contiguous match of 10 words or more therefore almost always requires verbatim reproduction of at least one full utterance, whereas a shorter threshold would have penalized the stock descriptive phrases (e.g., references to the overflowing sink, the boy on the stool) that recur across independent descriptions of the same picture regardless of memorization.

No retained narrative reached the 10-word criterion in any of the six cells. The longest contiguous overlap observed was 9 words in five of the six cells (Gemma-Dementia, Gemma-Healthy, Llama-Dementia, Llama-Healthy, Qwen-Healthy) and 8 words in the sixth (Qwen-Dementia), with per-cell median overlaps of 5–7 words and 90th-percentile overlaps of 6–9 words – consistent with shared stock phrasing across independent descriptions of the same picture rather than reproduction of training transcripts.

Supplementary Note 4 | WRAP versus Pitt: cohort and collection differences

**Demographics.** Pitt draws its Healthy/Dementia contrast from 99 controls (age 64.8 ± 7.8) and 141 probable-AD patients (71.6 ± 8.5); WRAP's held-out test set is older overall and less separated by group – 787 healthy controls (70.0 ± 7.3) versus 120 MCI (73.9 ± 6.0) – and, critically, the impaired class is MCI, a milder and earlier stage of impairment than the frank dementia the models were fine-tuned to distinguish, at a natural 13.2% (6.6:1) prevalence rather than Pitt's roughly balanced design.

Supplementary Note 5 | Pooled four-task narrative analysis

**Augmentation across narrative tasks.** The primary Augmentation analysis used synthetic Cookie Theft narratives, matching the elicitation task available in WRAP. We additionally tested whether incorporating synthetic narratives from all four generation tasks—Cookie Theft, Cinderella retelling, sandwich preparation and general memories—altered performance.

Using the same augmentation budget as the primary analysis (600 synthetic narratives), pooling across all four tasks yielded ROC-AUC = 0.574, PR–AUC = 0.177, MCI-F1 = 0.259 and MCI recall = 0.633 at the prespecified screening-oriented operating point. MCI-F1 remained significantly higher than the unaugmented baseline (0.211; $p = .044$), although ROC-AUC did not differ significantly from baseline. These results were similar, but slightly weaker, than augmentation using Cookie Theft narratives alone (ROC-AUC = 0.586, PR–AUC = 0.185, MCI-F1 = 0.268, recall = 0.717; $p = .020$ for MCI-F1 versus baseline).

We also evaluated an uncapped four-task pool containing all 2,345 available synthetic narratives. Performance was again similar to the primary Cookie-Theft-only analysis (ROC-AUC = 0.585, PR–AUC = 0.189, MCI-F1 = 0.266 and recall = 0.650), with MCI-F1 remaining significantly higher than baseline ($p = .035$). PR–AUC was not separately subjected to paired significance testing in these analyses.

Thus, broadening the synthetic augmentation set across narrative tasks did not improve performance over Cookie Theft augmentation alone. At a matched augmentation budget, performance was modestly lower, whereas increasing the size of the four-task pool approximately recovered the performance of the

Cookie-Theft-only condition. The overall conclusion—that dementia-conditioned synthetic language can improve minority-class prediction in WRAP—was therefore not specific to the use of Cookie Theft generations.

Supplementary Note 6 | Iowa Gambling Task payoff schedule, implementation, and behavioral features

The task follows the Experiment 1 payoff structure in Zemla & Davis (2025)[13]. Participants and LLM agents alike begin with a starting balance of $2,000.

Each deck carries its own fixed sequence of (win, loss) outcomes for its 1st through 40th draw. If a deck is drawn more than 40 times in a session, its payout resumes from the start of the sequence – the position indexed is (number of times that deck has been drawn) mod 40. The schedule is deterministic and identical across all agents (as it was across all human participants in the original study); no randomization is applied to deck order. The full deck-by-deck schedule is given in Supplementary Table 6.

**Behavioral features.** We represented each session using 50 features spanning eight groups: per-deck value and recency, per-deck loss statistics, good-vs-bad deck summaries, sequential learning signals, punishment sensitivity, loss-frequency/magnitude preference, running session history, and the current trial's outcome. Trial-level features were averaged over the session to form the classifier input. Undefined early-trial statistics were assigned predefined default values: 0.0/0 for count- and magnitude-based features and 0.5 for punishment-sensitivity and frequency-preference rates, representing a neutral prior. Post-loss and post-win punishment-sensitivity features remained at their default values until at least 10 trials of history were available. The complete feature list, formal definitions, and default values are provided in Supplementary Data 4.

Supplementary Note 7 | IGT prompt-configuration comparison and invalid-choice rates

We evaluated several prompt configurations for eliciting Iowa Gambling Task choices from each model family before the downstream synthetic-to-human analyses. Configurations included canonical deck-letter choices, shuffled color labels, and examiner-style prompting. Because model families differed in their responses to these formats, the elicitation configuration was selected separately for each family based on cross-validated Healthy–Dementia separability.

**Separability by configuration (5-fold cross-validated ROC-AUC, synthetic Healthy vs. Dementia).** For Gemma, canonical deck-letter prompting was selected (0.999 ± 0.001), compared with shuffled color labels (0.688 ± 0.020) and examiner-style prompting (0.965 ± 0.012). For Llama, shuffled color labels were selected (0.657 ± 0.014); performance was similar with examiner-style prompting (0.659 ± 0.027) and lower with deck-letter prompting (0.597 ± 0.037). For Qwen, examiner-style prompting was selected (0.803 ± 0.025), compared with canonical deck-letter prompting (0.633 ± 0.025) and shuffled color labels (0.629 ± 0.026).

**Invalid-choice rate.** Invalid choices were uncommon across configurations and model conditions, occurring on at most 3.3% of trials and on 0% of trials for most model–condition pairs.

Supplementary Note 8 | Breakdown-detection criteria for weight-space extrapolation

Generation breakdown was assessed automatically using a composite criterion, degenerate = repeat_flag | loop_flag | term_flag | refusal_flag, combining four checks.

**Repetition rate.** A narrative was flagged when its maximum 4-gram repetition rate exceeded 0.15, restricted to narratives of at least 20 words. Below 20 words a single repeated 4-gram can trivially score 1.0 in an otherwise unremarkable response, so shorter narratives were excluded from this check.

**Degenerate n-gram loop.** A narrative was flagged when its longest repeated-token run exceeded a family- and task-specific cutoff, defined as the 95th percentile of longest_run computed on the α=0 anchor data for that (family, task) pair, floored at 6 repeated tokens.
**Failure to terminate.** A narrative was flagged when generation reached the 200-token cap without ending on terminal punctuation.

**Generic AI-assistant language.** A narrative was flagged when its first 160 characters (REFUSAL_WINDOW_CHARS = 160) matched a case-insensitive regular expression covering assistant self-identification, inability statements, and refusal formulations:

```
_REFUSAL_RE = re.compile(
    r"as an? .{0,20}(artificial intelligence|\bai\b)( language model)?|"
    r"i'?m (just )?an? .{0,20}(artificial intelligence|\bai\b)|"
    r"as an? language model|"
    r"i don'?t have (the )?(ability|capability|access) to|"
    r"i (do not|don'?t) have access to|"
    r"my (abilit(y|ies)|functions?) (is|are) limited|"
    r"i (would not|wouldn'?t|am not able to|am unable to|cannot|can'?t) be able to|"
    r"i'?m sorry,? (but )?i (can'?t|cannot)",
    re.IGNORECASE,
)
```

**Breakdown-α determination.** For each family and direction, the breakdown threshold was defined as the baseline degenerate rate over α∈[0,1] plus max(0.10, 3×SE). The breakdown α was the smallest tested α ≥ 1.0 at which the degenerate rate crossed this threshold and remained above it at every larger tested α; a single point crossing the threshold without being sustained at larger α did not count. Families and directions not meeting this criterion within the tested range were right-censored as ‘no breakdown detected within the tested extrapolation range’.

#### Supplementary Note 9 | Linguistic-measure missing values and filled-pause lexicon

**Missing-value rules.** MATTR undefined for texts shorter than the window; repetition intervals undefined when nothing recurs; ratio measures undefined at zero denominators; Honoré undefined when all types are hapax. Undefined values are stored as missing, never imputed at computation time. Empirically, missingness in synthetic generations is low and slightly higher in the Dementia condition (e.g., bigram repetition interval 13.8% Dementia vs 11.2% Healthy; MATTR 5.7% vs 3.5%; all other measures ≤2.5% in both conditions, pooled across families/tasks); no measure was undefined for an entire condition in any family × task cell.

**Filled-pause lexicon** (verbatim, 21 items): um, uh, erm, er, ah, hmm, hm, mm, mhm, uh-huh, uhh, umm, ehm, eh, uhhh, ummm, huh, errm, eeh, ooh, aah.

#### Supplementary Data

##### Supplementary Data 1 (separate file)

Complete evaluation set for the instruction-following/constraint-satisfaction probe (n = 300 prompts, 7 categories)

##### Supplementary Data 2 (separate file)

Narrative-elicitation prompt bank (10 formulations × 4 tasks)

##### Supplementary Data 3 (separate file)

Iowa Gambling Task elicitation prompts, protocol comparison, and invalid-choice rates

##### Supplementary Data 4 (separate file)

Definitions of the 50 IGT trial-level behavioral features

##### Supplementary Data 5 (separate file)

Computed 50-feature behavioral vectors for real and synthetic IGT sessions

##### Supplementary Data 6 (separate file)

Clean/corrupted factual question-answer pairs for causal activation patching (n = 50)
